# Development of Gemcitabine-Modified miRNA Mimics as Cancer Therapeutics for Pancreatic Ductal Adenocarcinoma

**DOI:** 10.1101/2023.08.14.553255

**Authors:** John G. Yuen, Ga-Ram Hwang, Andrew Fesler, Erick Intriago, Amartya Pal, Anushka Ojha, Jingfang Ju

## Abstract

Pancreatic cancer, including its most common subtype, pancreatic adenocarcinoma (PDAC), has the lowest five-year survival rate among patients with pancreatic cancer in the United States. Despite advancements in anticancer treatment, the overall median survival for patients with PDAC has not dramatically improved. Therefore, there is an urgent need to develop new strategies of treatment to address this issue. Non-coding RNAs, including microRNAs (miRNAs), have been found to have major roles in carcinogenesis and the subsequent treatment of various cancer types like PDAC. In this study, we developed a treatment strategy by modifying tumor suppressor miRNAs, *hsa-miRNA-15a* (miR-15a) and *hsa-miRNA-194-1* (miR-194), with the nucleoside analog chemotherapeutic gemcitabine (Gem) to create Gem-modified mimics of miR-15a (Gem-miR-15a) and miR-194 (Gem-miR-194). In a panel of PDAC cell lines, we found that Gem-miR-15a and Gem-miR-194 induce cell cycle arrest and apoptosis, and these mimics are potent inhibitors with IC_50_ values up to several hundred fold less than their native counterparts or Gem alone. Furthermore, we found that Gem-miR-15a and Gem-miR-194 retained miRNA function by downregulating the expression of several key targets including WEE1, CHK1, BMI1, and YAP1 for Gem-miR-15a, and FOXA1 for Gem-miR-194. We also found that our Gem-modified miRNA mimics exhibit an enhanced efficacy compared to Gem alone in patient-derived PDAC organoids. Furthermore, we observed that Gem-miR-15a significantly inhibits PDAC tumor growth *in vivo* without observing any noticeable signs of toxicity. Overall, our results demonstrate the therapeutic potential of Gem-modified miRNAs as a treatment strategy for PDAC.

**One Sentence Summary:** Yuen and Hwang *et. al.* have developed a potent therapeutic strategy for patients with pancreatic cancer by modifying microRNAs with gemcitabine.

## INTRODUCTION

Pancreatic cancer, including the most common subtype pancreatic ductal adenocarcinoma (PDAC), is the 4^th^ leading cause of cancer-related death in the United States and has the lowest five-year relative survival rate, at 12%, despite being the 8^th^ and 10^th^ most common type of cancer in women and men, respectively (*1, 2*). Due to its difficulty in early detection, most patients present with advanced disease with an intrinsic resistance to chemotherapy (*3, 4*). Resistance to chemotherapy is common due to the complex biology of PDAC, largely mediated by the tumor microenvironment (TME) of pancreatic cancer (*5*). Therefore, combination chemotherapy regimens, gemcitabine (Gem) with albumin-bound paclitaxel or FOLFIRINOX (5-fluorouracil, irinotecan, leucovorin, and oxaliplatin), are the standard therapy to overcome this intrinsic resistance (*2, 6–8*). However, these regimens are still largely ineffective, as the median overall survival is only marginally improved relative to monotherapy, and the overall survival has not dramatically improved in the past 30 years (*9*). Despite advancements in pancreatic cancer research in both early detection and treatment, its limited impact on patient outcomes reveals an urgent need for novel therapies.

MicroRNAs (miRNAs) are a family of small, ∼22 nucleotide long, non-coding RNAs that are involved in RNA interference (RNAi), through the degradation of mRNA and/or translational repression, by binding to the 3’ UTR of their targets (*10*). This interaction is mediated by a 6-8-mer seed sequence located at the 5’ end of the miRNA, resulting in the regulation of multiple mRNA targets per miRNA. Dysregulation of miRNA expression results in various diseases, including cancer (*11, 12*). The therapeutic efficacy of miRNAs has been explored by restoring the expression of downregulated tumor suppressor miRNAs and by inhibiting overexpressed oncogenic miRNAs (*12–15*). However, despite the therapeutic potential of miRNA-based cancer therapeutics, clinically approved miRNA-based cancer therapeutics are in development, with none currently approved (*16, 17*).

*Hsa-miR-15a* (miR-15a) and *hsa-miR-194-1* (miR-194) are dysregulated in PDAC, and our lab has shown that low miR-15a expression correlates with poor prognoses patients in The Cancer Genome Atlas (TCGA) Pan Cancer Atlas, suggesting an important role for miR-15a in PDAC.(*18–20*) Meanwhile, downregulated expression of miR-194 has been previously reported in PDAC samples (*20*). Specifically, a correlation between low expression of miR-194 and the basal-like subtype, an aggressive subtype of PDAC, was observed (*20–23*). We discovered that miR-15a suppresses several key elevated oncogenic targets in PDAC including, polycomb complex protein BMI-1 (BMI1), checkpoint kinase 1 (CHK1), WEE1 G2 checkpoint kinase (WEE1), and Yes1 associated transcriptional regulator (YAP1) (*18*). BMI1 is a commonly overexpressed cancer stem cell marker, associated with poor prognosis and plays many roles in PDAC, including regulating proliferation, self-renewal, epithelial-to-mesenchymal transition (EMT), and metastasis (*24, 25*). YAP1 is a transcriptional co-activator, activating oncogene expression downstream of the Hippo signaling pathway and promotes the bypass of oncogenic KRAS addiction, directly contributing to oncogenesis in PDAC (*25–27*). Both WEE1 and CHK1 are key G2 checkpoint kinases that have been recently evaluated as targets in clinical trials for treating PDAC (*28–31*). Meanwhile, miR-194 has also been reported to target expression of BMI1 in endometrial cancer and targets forkhead box protein A1 (FOXA1) in lung cancer (*32, 33*). Dysregulation of FOXA1 expression has been shown to affect apoptosis and invasion in several cancer types, including lung cancer, liver cancer, and PDAC (*33–35*). Although these targets are important in its biology, the survival benefit of singularly targeting some of these proteins, such as WEE1, in patients with PDAC is minimal (*30, 31*). Combined with the overall poor response to traditional chemotherapeutics, the complex nature of resistance and the limitations of a single-target approach has been a major obstacle in the development of cancer therapeutics for patients with PDAC (*4, 36*). Therefore, the pleiotropic nature of tumor suppressor miRNAs such as miR-15a and miR-194 may hold therapeutic potential in overcoming this obstacle.

We have begun to address this gap in knowledge, demonstrating that certain tumor suppressor miRNAs—miR-15a and miR-129—are downregulated in colon cancer and regulates key oncogenes, including Bcl-2, BMI1, YAP1, and DCLK1 (*37, 38*). We further show that miRNA mimics—created by replacing the uracils (U) on the guide strand of the miRNA with 5-fluorouracil (5-FU), a nucleoside analog antimetabolite chemotherapeutic—are highly effective at inhibiting colon, lung, and pancreatic cancer cells both *in vitro* and *in viv*o (*18, 37–39*). Notably, these miRNA mimics retain specificity to their mRNA targets, exert chemotherapeutic effects through its modification, and gain the ability to be delivered to cancer cells without the aid of a delivery vehicle, a major bottleneck for nucleic acid-based drug development. In this study, we developed a different miRNA modification strategy by using Gem to create a more potent miRNA mimic for the treatment of PDAC. Using this strategy, we have developed Gem-modified miRNA mimics of miR-15a (Gem-miR-15a) and miR-194 (Gem-miR-194). Our study demonstrates the viability of our Gem modification strategy, as seen by the efficacy of Gem-miR-15a and Gem-miR-194 at inhibiting cell proliferation and inducing cell cycle arrest and apoptosis *in vitro* with and without the use of a delivery vehicle. Our Gem-modified miRNA mimics also retain function as miRNAs, and notably, our results demonstrate the potential for Gem-miR-15a as a potent cancer therapeutic for PDAC *in vivo*. This study establishes a platform strategy for modifying miRNAs as multi-targeted cancer therapeutics to potentially improve current anti-cancer therapies.

## RESULTS

### Development of Gemcitabine-Modified miRNA Mimics

To evaluate the therapeutic potential of combining both miRNA-based therapeutics and gemcitabine (Gem), we designed and developed gemcitabine-modified miRNA mimics. By substituting the endogenous cytidine (C) bases of miR-15a and miR-194 with Gem, we developed the gemcitabine-modified mimics, Gem-miR-15a and Gem-miR-194 (Fig. 1A). Notably, we only modified the guide strands of these mimics, and the passenger strands were left unmodified to avoid potential off-target effects and to preserve miRNA function.

**Fig. 1.**
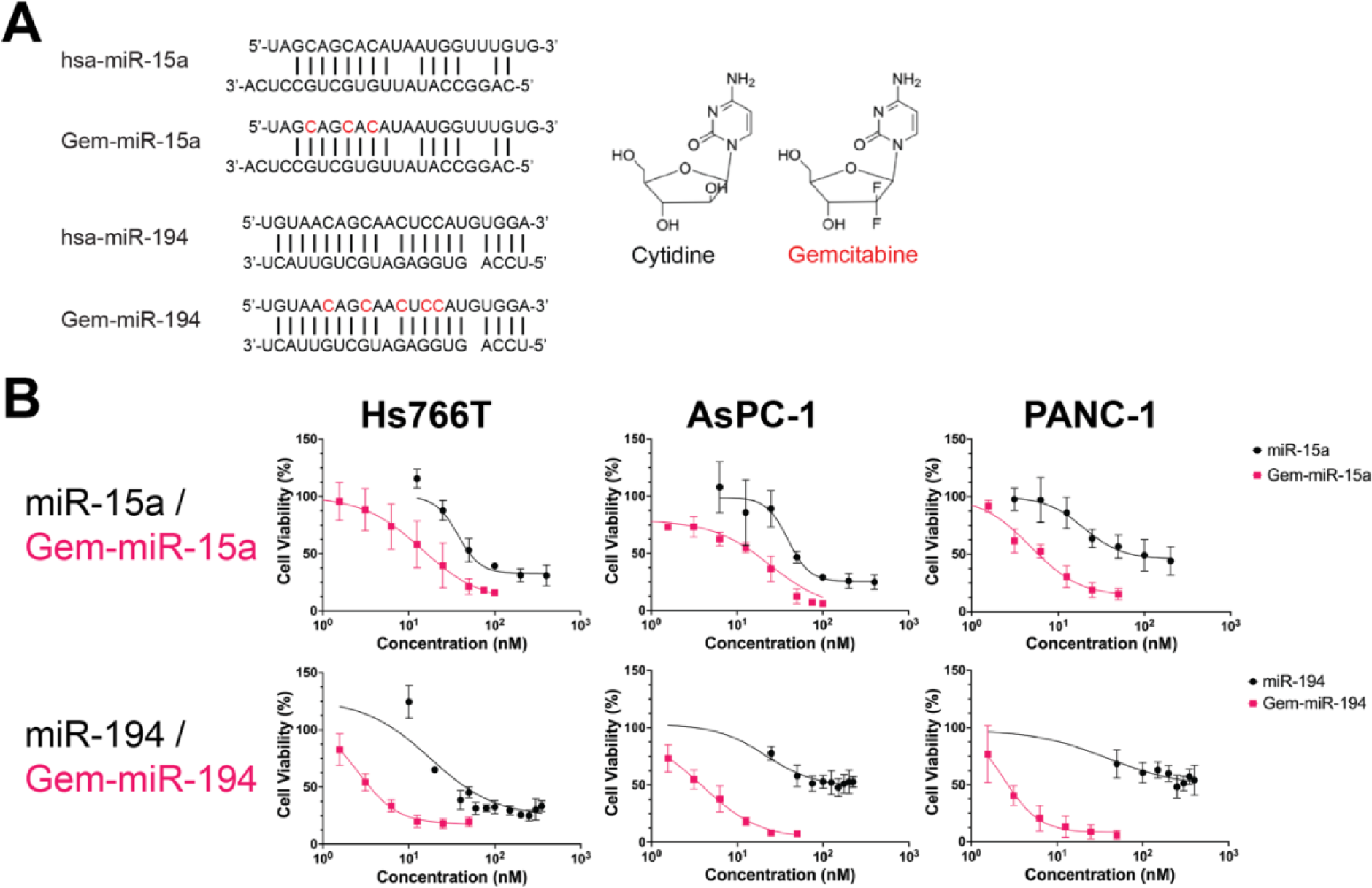
Gem-modified miRNA mimics exhibit enhanced inhibition of cell proliferation compared to their unmodified counterparts in PDAC *in vitro* without the use of a transfection vehicle. **(A)** Gem-modified miRNA mimics of miRNAs miR-15a (Gem-miR-15a) and miR-194 (Gem-miR-194) were developed by substituting cytidine within their respective guide strands with the cytidine nucleoside analog, gemcitabine (red). **(B)** Compared to their unmodified counterparts, Gem-miR-15a and Gem-miR-194 were found to be more effective at inhibiting cell proliferation in PDAC cell lines Hs766T, AsPC-1, and PANC-1 (n = 4). Gem-modified mimics were administered to PDAC cells through vehicle-free delivery. IC_50_ values of these treatment conditions are listed in Table 1. Data are represented as mean ± SD.

**Table 1.** IC_50_ values of Gem-miR-15a, Gem-miR-194 and Gem in various 2D PDAC cell lines and hTERT-HPNE cells. Data are represented as mean ± SD.

|  | miR-15a | Gem-miR-15a | miR-194 | Gem-miR-194 | Gemcitabine |
| --- | --- | --- | --- | --- | --- |
| *Hs766T* | 51.46 ± 10.83 | 16.23 ± 8.30 | 25.33 ± 11.69 | 3.56 ± 1.52 | 8,677.60 ± 2,679.29 |
| *AsPC-1* | 47.96 ± 11.94 | 15.94 ± 20.22 | >200 nM | 3.79 ± 3.41 | >10,000 nM |
| *PANC-1* | 70.63 ± 11.68 | 5.86 ± 1.64 | >200 nM | 2.68 ± 2.61 | 6,143.23 ± 1,575.14 |
| *MIA PaCa-2* | | 9.86 ± 0.96 | | | |
| *Capan-1* | | 8.34 ± 1.21 | | | |
| *HPNE* | >200 nM | 4.77 ± 9.88 | | | |

### Gem-miR-15a and Gem-miR-194 Display Enhanced Efficacy at Inhibiting PDAC Proliferation *In Vitro* Without Use of a Delivery Vehicle

To demonstrate that Gem-miR-15a and Gem-miR-194 display an enhanced efficacy at inhibiting cancer proliferation *in vitro*, using a WST-1 dye proliferation assay, we performed a dose response assay to determine the IC_50_ values of these mimics in three different PDAC cell lines, Hs766T, AsPC-1 and PANC-1 (Fig. 1A & Table 1). Compared to their native counterparts, miR-15a and miR-194, we observed a 3.2 to 12.1-fold change difference in the IC_50_ value for Gem-miR-15a and a 7.1 to 74.6-fold change difference in the IC_50_ value for Gem-miR-194. With these three cell lines, we also performed a dose response assay for Gem, and we observed a 534.7 to 1,048.3-fold change difference in the IC_50_ value between Gem and Gem-miR-15a and a 2,292.3 to 2,638.5-fold change difference between Gem and Gem-miR-194 (Table 1). It is important to note that for Gem-miR-15a and Gem-miR-194, these assays were performed without the use of a delivery vehicle to also demonstrate cellular uptake of these Gem-modified miRNA mimics in PDAC cells. However, a delivery vehicle was used with miR-15a and miR-194 as miR-15a as unmodified miRNAs cannot typically cross the lipid bilayer of the cellular membrane of cells due to their negative charges.

In addition to the previously mentioned PDAC cell lines, we also examined the efficacy of Gem-miR-15a in PDAC cell lines, MIA PaCA-2 and Capan-1, and we observed IC_50_ values similar to the other PDAC cell lines (IC_50_ = 9.86 ± 0.96 nM and 8.34 ± 1.21 nM, respectively) (Table 1). We also examined and compared the efficacies of miR-15a and Gem-miR-15a in the immortalized pancreatic epithelial cell line, hTERT-HPNE. We observed that hTERT-HPNE was generally insensitive to miR-15a (IC_50_ = >200 nM) while being sensitive to Gem-miR-15a (IC_50_ = 4.77 ± 9.88 nM). Overall, our results suggest that our Gem-modified miRNAs effectively inhibit cell proliferation and can be delivered without the use of a delivery vehicle.

### Gem-miR-15a and Gem-miR-194 Induces Cell Cycle Arrest and Apoptosis in PDAC

To explore the potential mechanisms by which cancer cell proliferation is inhibited, we assessed the effects of our Gem-modified miRNA mimics on cell cycle. In the PDAC cell line Hs766T, we observed a significant increase of cells in G1 phase after treatment with Gem-miR-15a and Gem-miR-194 compared to cells treated with a non-specific miRNA mimic (negative control) (Fig. 2A & 2B). In addition, we observed a significant decrease of cells in S phase after treatment with Gem-miR-15a and a significant decrease of cells in G2 phase after treatment with Gem-miR-194. Our results also show a 5.3-fold and 1.9-fold change in G1/S ratio (*p* = 0.0002 and *p* = 0.0315) and a 0.4-fold and 0.8-fold change in G2/G1 ratio (*p* = 0.0463 and *p* < 0.0001) after treatment with Gem-miR-15a and Gem-miR-194, respectively. In addition to cell line Hs766T, we also assessed the effects of Gem-miR-15a and Gem-miR-194 in the PDAC cell lines, AsPC-1 and PANC-1 (Fig. S1). In AsPC-1 cells, our results show a significant increase in G1/S ratio (2.6-fold change, *p* < 0.0001 and 2.6-fold change, *p* < 0.0001) and a significant decrease in G2/G1 ratio (< 0.1-fold change, *p* < 0.0001 and < 0.1-fold change, *p* < 0.0001) after treatment with Gem-miR-15a and Gem-miR-194, respectively. In PANC-1 cells, we only observed a significant increase in G1/S ratio after treatment with Gem-miR-194 (1.9-fold change, *p* = 0.0030). However, we also observed a significant decrease in G2/G1 ratio after treatment with Gem-miR-15a (0.9-fold change, *p* = 0.0001) and a significant increase in G2/G1 ratio after treatment with Gem-miR-194 (1.5-fold change, *p* = 0.0090). Overall, our results suggest that our Gem-modified miRNAs induce cell cycle arrest, but depending on the cell line and miRNA mimic, it can be either at the G1 phase (Hs766T and AsPC-1 cells) or at the G2 phase (PANC-1 cells with Gem-miR-194).

**Fig. 2.**
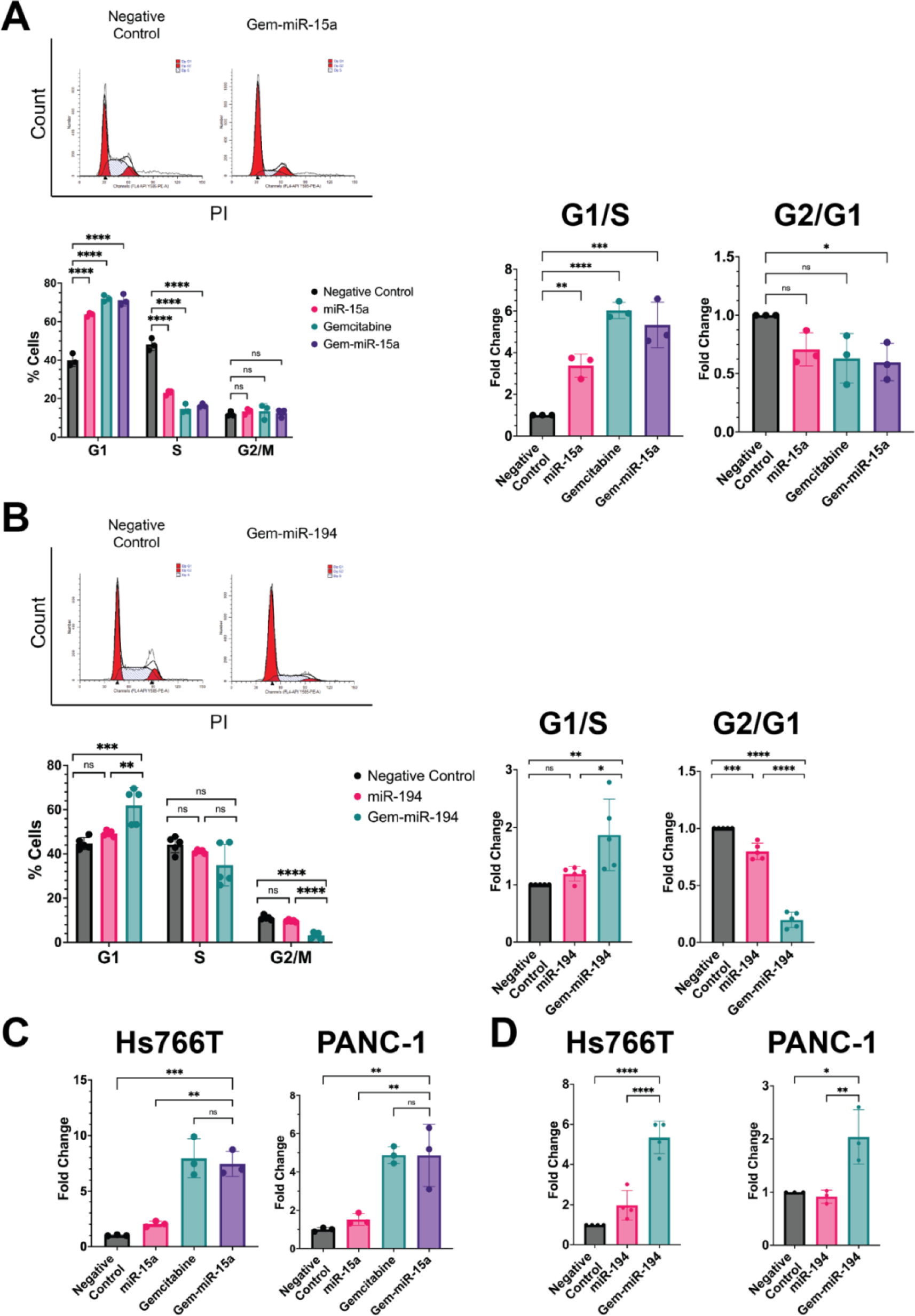
Gem-miR-15a and Gem-miR-194 induce cell cycle arrest and apoptosis in PDAC *in vitro*. **(A)** Gem-miR-15a was found to induce G1 cell cycle arrest in the PDAC cell line Hs766T, as seen by an increase of cells in G1 (*p* < 0.0001) and a decrease in S phase (*p <* 0.0001) (n = 3). G1/S (*p* = 0.0002) and G2/G1 (*p* = 0.0463) ratios were also calculated, suggesting G1 arrest. **(B)** Gem-miR-194 was also found to induce G1 cell cycle arrest, as seen by an increase of cells in G1 (*p* = 0.0004) and a decrease of cells in G2 (*p* < 0.0001) (n = 5). G1/S (*p* = 0.0073) and G2/G1 (*p* < 0.0001) ratios were also calculated, suggesting G1 arrest. **(C)** Gem-miR-15a and **(D)** Gem-miR-194 were also found to induce apoptosis in PDAC cell lines Hs766T (*p* = 0.0003, n = 3 & *p* < 0.0001, n = 4) and PANC-1 (*p* = 0.0025, n = 3 & *p* = 0.0134, n = 4). Data are represented as mean ± SD. \**p* < 0.05, \*\**p* < 0.01, \*\*\**p* < 0.001, \*\*\*\**p* < 0.0001.

In addition to cell cycle, we also assessed the effects of our Gem-modified miRNA mimics on apoptosis. We observed that Gem-miR-15a induces apoptosis in Hs766T cells (7.5-fold increase, *p* = 0.0003) and PANC-1 cells (4.9-fold increase, *p* = 0.0025) (Fig. 2C). Comparatively, miR-15a does not significantly induce apoptosis, but treatment with Gem alone causes an 8.0 and 4.9-fold change increase in apoptosis in Hs766T and PANC-1 cells, respectively (*p* = 0.0002 and *p* = 0.0024). Likewise, we observed that Gem-miR-194 also induces apoptosis in Hs766T cells (5.3-fold increase, *p* < 0.0001) and PANC-1 cells (2.0-fold increase, *p* = 0.0143). Therefore, our results demonstrate that our Gem-modified miRNA mimics also induce apoptosis in PDAC cells.

### Gem-miR-15a and Gem-miR-194 Retain Target Specificity in PDAC

To demonstrate that Gem-miR-15a acts as a miRNA mimic–retaining its ability to induce RNAi– we investigated its ability to downregulate previously reported targets of miR-15a, BMI1, CHK1, WEE1, and YAP1. In PANC-1, AsPC-1, and Hs766T cells, Gem-miR-15a was found to decrease BMI1, CHK1, and WEE1 expression (Fig. 3A). Similarly, Gem-miR-15a decreased YAP1 expression in PANC-1 and AsPC-1 cells. Notably, because these targets have been previously reported to be downregulated by miR-15a and a miR-15a miRNA mimic modified with the nucleoside analog 5-FU in PDAC, we treated these cells with Gem-miR-15a without the use of a transfection vehicle to also demonstrate that Gem-miR-15a can downregulate these targets without the use of a transfection vehicle (*18*).

**Fig. 3.**
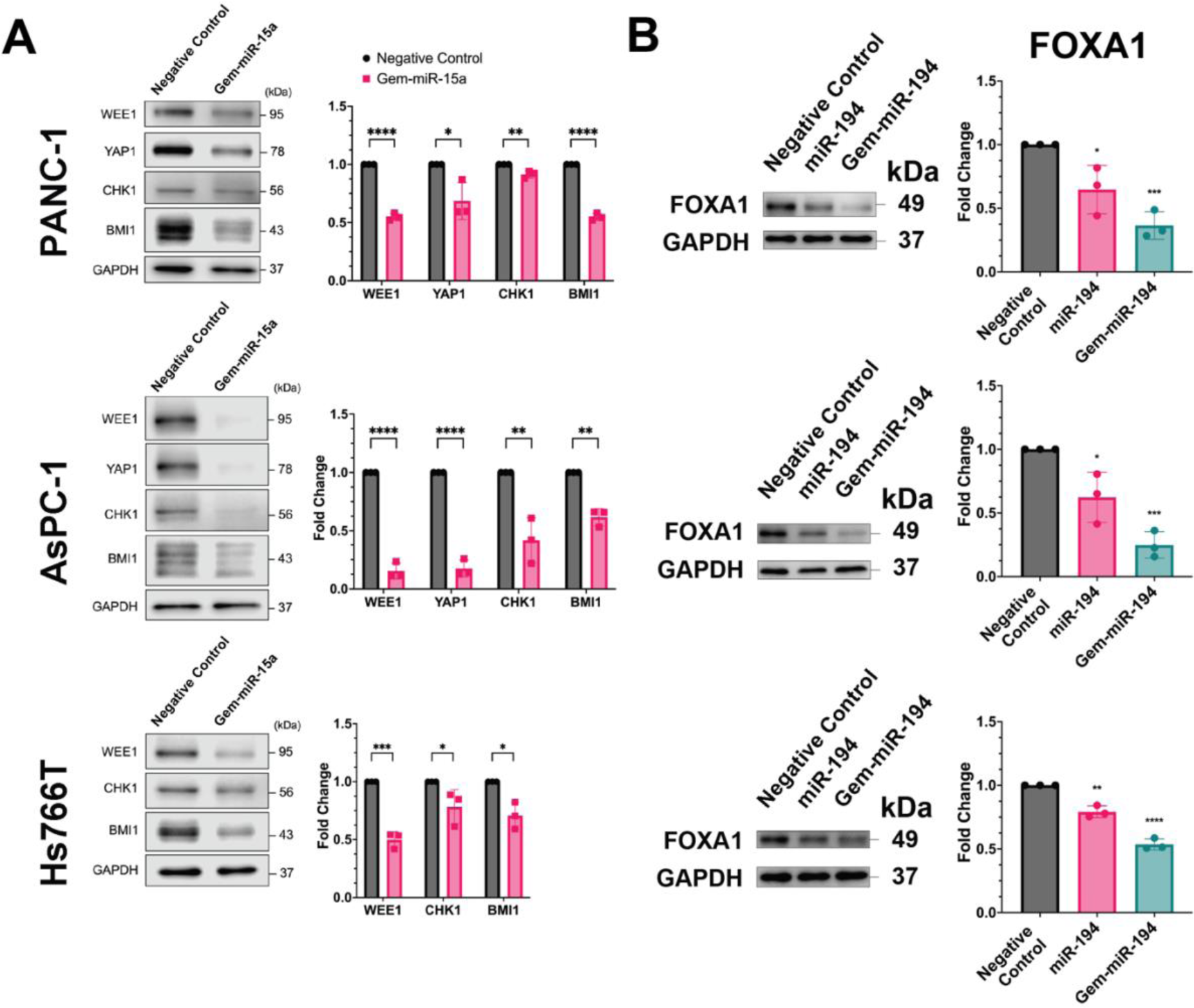
Gem-miR-15a and Gem-miR-194 retain miRNA function by downregulating expression of respective targets with and without the use of a delivery vehicle. **(A)** Gem-miR-15a was found to retain miRNA function by downregulating miR-15a targets WEE1, CHK1, and BMI1 in PDAC cell lines PANC-1, AsPC-1 and Hs766T (n = 3). Gem-miR-15a was also found to downregulate expression of miR-15a target YAP1 in PDAC cell lines PANC-1 and AsPC-1 (n = 3). PDAC cell lines were transfected with Gem-miR-15a through vehicle-free delivery. **(B)** Gem-miR-194 and miR-194 were found to downregulate expression of miR-194 targets FOXA1 in PDAC cell lines Hs766T, AsPC-1, and PANC-1 (n = 3). PDAC cell lines were transfected with Gem-miR-194 with the use of a delivery vehicle. Data are represented as mean ± SD. \**p* < 0.05, \*\**p* < 0.01, \*\*\**p* < 0.001, \*\*\*\**p* < 0.0001.

Because miR-194 has been reported to downregulate different targets from miR-15a, we investigated Gem-miR-194’s ability to downregulate targets FOXA1 and BMI1 (Fig. 3B & S2). Gem-miR-194 was found to decrease FOXA1 and BMI1 expression in PANC-1 and Hs766T cells. Gem-miR-194 was also found to decrease FOXA1 expression in AsPC-1 cells (Fig. 3B). However, miR-194 was found to decrease only FOXA1 in these PDAC cell lines, and miR-194 was not found to downregulate BMI1 expression (Fig. S2).

We also examined whether our results were due to off-targeting effects acquired by incorporating Gem into our miRNA mimics. We developed a Gem-modified non-specific control miRNA mimic by modifying a non-specific control miRNA sequence, *cel-miR-67* (cel-67) with Gem (Gem-cel-67). Using CHK1 as a target, we investigated Gem-cel-67’s ability to downregulate CHK1 expression in PANC-1, and we did not observe a significant change in CHK1 expression (Fig. S3), suggesting that our Gem-modified miRNAs retain their function as miRNA mimics.

### Gem-miR-15a Is Loaded into the RISC in PDAC Under Vehicle-Free Conditions

To further show that our Gem-modified miRNA mimics retain miRNA behavior and interact directly with RNAi machinery, we performed AGO-crosslinking and immunoprecipitation (AGO-CLIP) (*40*). Using Gem-miR-15a, we used AGO-CLIP to see whether Gem-miR-15a is loaded into AGO and can recruit its target mRNAs in PANC-1 cells, specifically *CHK1* and *WEE1* (Fig. 5). Through RT-PCR of samples treated Gem-miR-15a, miR-15a was found to be enriched by 15386.8-fold, *CHK1* mRNA enriched 24.2-fold, and *WEE1* mRNA enriched 26.0-fold relative to treatment with negative control. This enrichment demonstrates that both the GEM-miR-15a and its targets, *CHK1* and *WEE1*, are loaded into AGO and the RNA-induced silencing complex (RISC). Notably, like in our dose response assays, transfection with Gem-miR-15a was performed without the use of a delivery vehicle.

### Gem-miR-15a and Gem-miR-194 Inhibit the Growth of PDAC Organoids and Gemcitabine Resistant Organoids

To demonstrate the efficacy of our Gem-modified miRNA mimics beyond two-dimensional PDAC cell lines, we examined the efficacy of Gem-miR-15a and Gem-miR-194 *ex vivo* using patient derived organoid models of PDAC (*41*). Three patient-derived organoid lines were selected to represent the basal-like (hF3) and classical (hF44, hT89) molecular subtype. Using these three organoid lines, we first determined the IC_50_ values of these lines to Gem with IC_50_ values ranging from 416.52 ± 287.06 nM to 508.89 ± 93.70 nM (hT89 and hF44, respectively) (Table 2). We then calculated the IC_50_ values of Gem-miR-15a on these three lines, and we observed that these organoids were more sensitive to Gem-miR-15a with a 145, 170, and 43-fold difference in IC_50_ values (hF44, hF3, and hT89, respectively) (Fig. 5 & Table 2). In addition to Gem-miR-15a, we also examined the efficacy of Gem-miR-194 in the organoid lines hF44 and hF3 (Fig. 5 & Table 2). Compared to Gem, we observed a 150 and 164-fold difference in IC_50_ values for Gem-miR-194 in hF44 and hF3, respectively. Overall, our results suggest that Gem-miR-15a and Gem-miR-194 are also effective at inhibiting PDAC organoid growth.

**Table 2.** IC_50_ values of Gem-miR-15a, Gem-miR-194 and Gem in various lines of PDAC organoids. Data are represented as mean ± SD.

|  | Gem-miR-15a | Gem-miR-194 | Gemcitabine |
| --- | --- | --- | --- |
| *hF44* | 3.50 ± 1.42 | 3.38 ± 3.00 | 508.89 ± 93.70 |
| *hF3* | 2.60 ± 0.70 | 2.70 ± 1.45 | 442.98 ± 43.16 |
| *hT89* | 9.54 ± 3.04 | | 416.52 ± 287.06 |

### Gem-miR-15a Inhibits PDAC Tumor Growth *In Vivo*

To demonstrate the therapeutic potential of Gem-miR-15a in PDAC *in vivo*, we investigated its effects on a metastatic PDAC mouse model. Tumor xenografts of luciferase-expressing Hs766T cells (Hs766T (+Luc)) in NOD-SCID mice were treated with either a vehicle control (vehicle), Gem, Gem-miR-194, or Gem-miR-15a, and a decrease in tumor growth was observed in mouse groups treated with either Gem or Gem-miR-15a via luciferase expression (Fig. 6). Compared to the vehicle control group, a 5.5-fold decrease was observed in the Gem-miR-15a (4 mg/kg) treated group (*p* = 0.0412), and a 13.0-fold decrease was observed in the Gem (50 mg/kg) treated group (*p* = 0.0083). In addition, body mass was measured and a rapid change in mass was calculated as an indicator of acute toxicity, which stayed within normal limits (<15% decrease in mass). However, a significant change in tumor growth was not observed in mice treated with Gem-mir-194 despite its potency in treating PDAC cell lines and patient-derived PDAC organoids.

Using the same metastatic PDAC mouse model, the effects of Gem-miR-15a *in vivo* was repeated to confirm the therapeutic potential of Gem-miR-15a (Fig. S4). In agreement with our previous results, a significant decrease in tumor growth was observed in mouse groups treated with either Gem-miR-15a (4 mg/kg) or Gem (12 mg/kg) (5.0-fold change decrease, *p* = 0.0028 and 3.4-fold change decrease, *p* = 0.0376, respectively). Body mass was also measured in these groups, and rapid changes in body mass were not observed.

## DISCUSSION

The poor response of patients with PDAC to therapies has been attributed to several mechanisms, including, intrinsic resistance from cancer stem cells, dysregulation of the cell cycle, and acquired resistance to anticancer treatment (*3–5*). In this study, we aimed to design an improved miRNA-based therapy to treat pancreatic cancer by modifying tumor suppressor miRNAs, miR-15a and miR-194, with Gem. In previous studies, dysregulation of miR-15a and miR-194 have been shown in pancreatic cancer, with their expressions associated with patient survival (*18–20, 42*). Furthermore, previous studies have also demonstrated the therapeutic potential of modifying tumor suppressor miRNAs with the nucleoside analog chemotherapeutic, 5-FU, in several cancers (*18, 37–39, 43, 44*). As seen by its ability to inhibit proliferation, induce cell cycle arrest and apoptosis in several different PDAC cell lines, our results demonstrate the therapeutic potential of Gem-miR-15a in PDAC and modifying miRNAs with Gem as a method of anticancer treatment. Compared to the IC_50_ values of their unmodified miRNA counterparts, Gem-miR-15 and Gem-miR-194 exhibit IC_50_ values that are several folds lower. Furthermore, Gem-miR-15a and Gem-miR-194 deliver Gem at doses hundreds of folds less than (∼760-fold and ∼2500-fold less, respectively) the IC_50_ value of Gem in PDAC cells, thus suggesting that these modified Gem-modified miRNAs sensitize PDAC cells to Gem. Therefore, our results demonstrate that modifying miRNAs with Gem is an effective strategy at sensitizing PDAC to cancer therapeutics.

In addition to 2D cell cultures, the advantages of testing the efficacy of cancer therapeutics in patient-derived 3D organoid cultures has been well-studied (*41, 45, 46*). These advantages include the high-throughput drug screening power that organoids provide compared to an *in vivo* model. With the development of patient-derived PDAC organoids, these organoids have been successfully utilized as an *ex vivo* platform to design a more personalized treatment regimen for PDAC (*41*). Due to these advantages, we also examined the efficacy of Gem-miR-15a and Gem-miR-194 in multiple patient-derived organoid lines of PDAC. Our results demonstrate the efficacy of both Gem-miR-15a and Gem-miR-194 in patient-derived organoid models of PDAC. In addition, compared to the IC_50_ values of Gem in these lines, it is worth noting that the IC_50_ values for both Gem-modified miRNAs in these organoid models were all < 10 nM and ranged from ∼40 to 170-fold more sensitive than the IC_50_ value of Gem. Consistent with previous studies, our data suggests that chemotherapy-modified miRNA-based therapies can improve therapeutic sensitivities by combining the power of chemotherapeutics and tumor suppressor miRNAs.

Dysregulation of miR-15a has been reported in PDAC, and our lab has previously reported a correlation between low expression of miR-15a and poor patient prognosis, thus suggesting an important role for miR-15a in PDAC (*18, 19*). The data also shows that despite being co-transcribed with miR-15, miR-15a is differentially processed during the maturation of pre-miR-15a to the mature miR-15a, as the mature miR-16 expression remains unchanged in PDAC (*18*). We also show that miR-15a suppresses several elevated oncogenic targets in PDAC including, BMI1, CHK1, WEE1, and YAP1. BMI1 is a marker that has been observed to be overexpressed in cancer stem cells, associated with poor prognosis, and plays many roles in PDAC, including regulating proliferation, self-renewal, and invasion (*24, 25*). The overexpression of BMI1 was observed in PanIN lesions, and PDAC cell lines and patient samples, and has been correlated with lymph node metastasis and poor survival (*24, 25, 47*). YAP1 is a transcriptional co-activator that is normally phosphorylated by active tumor suppressive Hippo signaling and sequestered into the cytoplasm (*48*). When Hippo signaling is inactive, YAP1 can drive the expression of genes associated with tissue healing, remodeling, and homeostasis. YAP1 has been found to be overexpressed in many types of cancer and is, therefore, considered an oncogene (*49, 50*). In PDAC, it was also found that YAP1 can substitute for oncogenic KRAS, signaling potentially contributing to resistance to therapy (*25, 26*). WEE1 and CHK1 are key G2/M checkpoint kinases that can affect CDK2 activation (*51, 52*). Both WEE1 and CHK1 have been recently evaluated in clinical trials for treating PDAC, and although they are important in its biology, the survival benefit of targeting WEE1 or CHK1 alone in patients with PDAC is minimal (*29–31*). However, our results demonstrate a correlation between the knockdown of these oncogenic targets and the induction of cell cycle arrest and apoptosis in PDAC cells (Fig. 2 & S1). Therefore, our results also demonstrate the advantages of utilizing a multi-targeted approach conferred by the inherent nature of miRNAs to better overcome resistance.

Dysregulation of miR-194 in several cancers, including PDAC, has also been reported in previous studies (*20, 33, 53, 54*). Specifically, miR-194 has been found to be downregulated in several cancers, including, a TCGA analysis demonstrating the correlation between the knockdown of miR-194 expression and the basal-like subtype of PDAC, aggressive subtype of PDAC known to be resistant to chemotherapy (*20–23*). This suggests that miR-194 is a tumor suppressor, thus making miR-194 a promising miRNA sequence to modify with Gem. In addition, targets of miR-194 include BMI1 and FOXA1 (*32, 33*). FOXA1 is a pioneer factor that binds to chromatin and facilitates the binding of additional transcription factors (*55*). Upregulation of FOXA1 has been found to inhibit apoptosis and promote metastasis in several cancers, including lung cancer, liver cancer, prostate cancer, and PDAC (*33–35, 55, 56*). Consistent with previous reports, our data shows that PDAC cells treated with Gem-miR-194 together with the downregulated expression of FOXA1 and BMI1, exhibited cell cycle arrest and apoptosis (*32–34*). However, despite having been previously reported as a target of miR-194 in endometrial cancer (*32*). miR-194 was not found have a significant impact on BMI1 expression in PDAC cells. This is highly consistent with previous studies that differential targets with the same miRNA may be largely due to different mRNA target abundance and while also being dependent on cellular context (*39, 57*). The suppression of Gem-miR-194 on BMI1 expression in PDAC may be due to the enhanced affinity binding to its mRNA target transcript compared to miR-194. Therefore, future studies utilizing RNA-seq to measure changes in mRNA expression after treatment with Gem-miR-194 may provide further insight into associated pathways, mechanisms, and changes in target affinity. However, overall, our results demonstrate, like Gem-miR-15a, Gem-miR-194 retains function as a miRNA by downregulating expression of targets of miR-194.

We have previously demonstrated in several cancer types, including colon, pancreatic, and breast cancer, that 5-FU modification of miRNAs not only allows for the preservation of miRNA function, but also confers its ability to enter the cell without the use of a delivery vehicle (*18, 37, 44*). Our results demonstrate that, like 5-FU modifications, Gem modification of miRNAs also grants the same cellular uptake property. Key oncogenic targets of miR-15a in PDAC, such as BMI1, CHK1, WEE1, and YAP1, can be downregulated without the use of a delivery vehicle. Furthermore, using AGO-CLIP, we also provide clear experimental evidence that Gem-miR-15a and its mRNA targets can be loaded onto AGO and can, therefore, participate in RNAi downstream. Combined with our previous observations of miR-15a regulating the expression of these targets in PDAC, this may provide additional evidence that the enhanced efficacy and deliverability of these modified miRNA mimics can be due to the increased lipophilicity that is conferred due to the addition of fluorine groups on the ribose ring of Gem to drug candidates, a strategy that has long been used for improving the lipophilicity of small molecule compounds (*18, 58*).

In addition to our *in vitro* data, our results demonstrate the promising potential of Gem-modified miRNAs as a cancer therapeutic within an *in vivo* context. Gem-miR-15a was found to significantly reduce PDAC tumor growth without having a significant impact on body weight, a metric used to monitor mouse health. It is important to note that our *in vivo* studies were performed in the immunodeficient NOD/SCID mouse model. Therefore, future studies within an immunocompetent mouse model can be used to assess Gem-miR-15a’s impact on a functional immune system.

Surprisingly, although Gem-miR-15a was found to significantly reduce tumor growth, Gem-miR-194 did not have a significant impact on PDAC tumor growth *in vivo* (Fig. S4). Although, initially, Gem-miR-194 seems to exert an inhibitory effect, however, a significant difference in PDAC tumor growth was not observed by the end. This is in contrast with our *in vitro* results that demonstrate Gem-miR-194’s potency as an inhibitor in both 2D cells and 3D organoid models (Fig. 1 & 4). This may reflect the complex cellular functions of one of Gem-miR-194’s targets, FOXA1, in cancer progression (*35, 55, 59*). Although a large body of evidence demonstrated that suppression of FOXA1 may be a good therapeutic strategy, however, the role of FOXA1 is rather complex (*55, 60, 61*). It has been also reported that FOXA1 mediated enhancer reprogramming increases metastatic potential (*35*). However, another study reported that direct inhibition of FOXA1 alone could induce EMT in PDAC to promote tumor progression (*59*). In breast cancer, downregulation of FOXA1 leads to cancer stem-like properties in tamoxifen-resistant breast cancer cells through induction of IL-6 (*62*). Meanwhile, expression of mutant FOXA1 has been found to promote prostate cancer invasion, and high expression of FOXA1 has been found to correlate with lower patient survival (*56, 63*). However, a separate study observed that loss of FOXA1 expression has also been to induce the TGF-β signaling pathway to induce EMT in prostate cancer, thus demonstrating the complex role of FOXA1 in cancer progression (*64*). Our results are also consistent with previous reports which observed that miR-194 initially inhibits PDAC tumor growth *in vivo*, followed by an accelerated growth of PDAC tumors resistant to radiation therapy (*65*). Therefore, our results also suggest the importance of including *in vivo* tumor model studies when developing cancer therapeutics. Our results also suggest that the impact of modifying tumor suppressor miRNAs with Gem is still dependent on the impact of each respective tumor suppressor miRNAs and their corresponding mRNA targets, thus emphasizing the importance of investigating the impact of a miRNA on a specific cancer type. For example, consistent, downregulated expression of miR-194 has been observed in colorectal cancer (CRC) and liver cancer (*54*). Therefore, future studies examining the therapeutic potential of Gem-miR-194 in these cancers may result in more promising results.

**Fig. 4.**
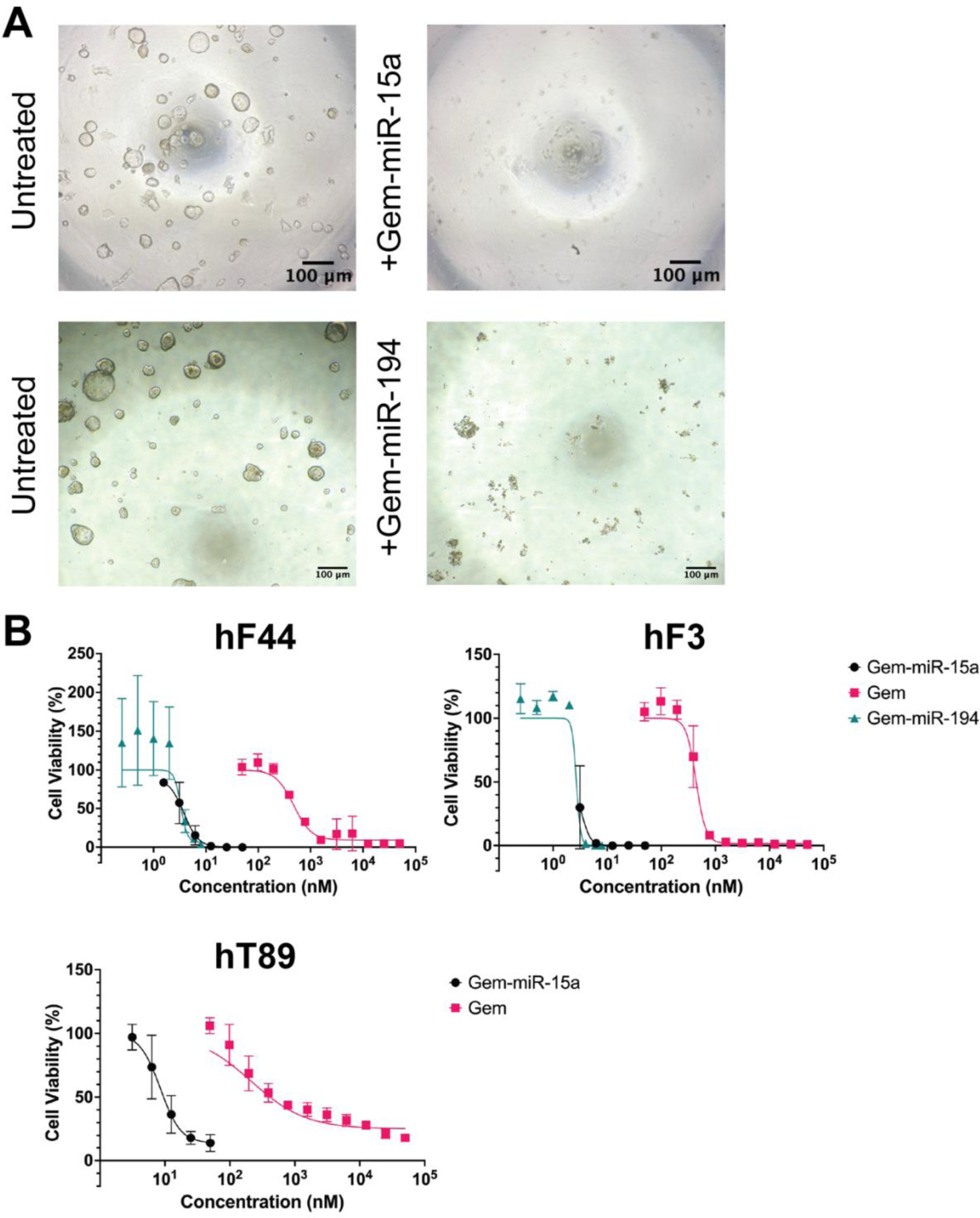
Gem-modified miRNA mimics exhibit enhanced inhibition of organoid growth in PDAC organoids without the use of a transfection vehicle. **(A)** Compared to the untreated group, growth of patient-derived PDAC organoids, hF44, was found to be inhibited by treatment with either Gem-miR-15a (12.5 nM) or Gem-miR-194 (8 nM). **(B)** Gem-miR-15a and Gem-miR-194 were found to be more effective at inhibiting organoid growth, as seen by dose response curves of Gem-miR-15a, Gem-miR-194 and Gem were plotted for PDAC organoids, hF44 and hF3 (n = 3). Compared to Gem, Gem-miR-15a was also found to be more effective at inhibiting organoid growth in PDAC organoid, hT89 (n = 3). IC_50_ values of Gem-miR-15a, Gem-miR-194 and Gem are listed in Table 2. Data are represented as mean ± SD.

**Fig. 5.**
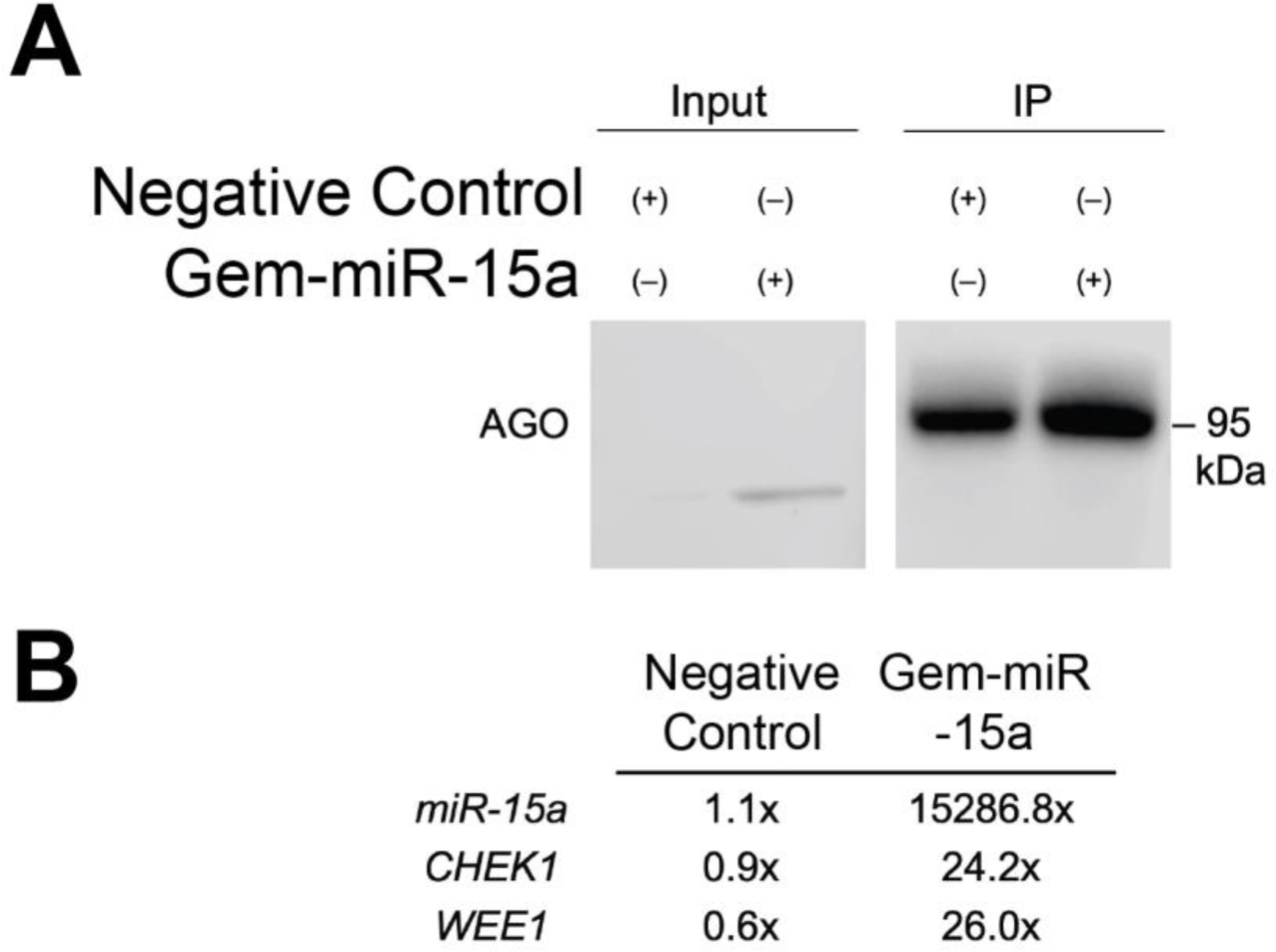
Gem-miR-15a and its targets are loaded into Argonaute. **(A)** PANC-1 cells were treated with either negative control or Gem-miR-15a, and cells were UV irradiated prior to immunoprecipitation of Argonaute (AGO-CLIP). **(B)** RNA was extracted from the same samples and RT-PCR was performed on miR-15a and two of its targets, *CHEK1* and *WEE1*. In Gem-miR-15a-treated AGO-CLIP samples, miR-15a was enriched by 15386.8-fold, *CHEK1* mRNA was enriched 24.2-fold, and *WEE1* mRNA was enriched 26.0-fold relative to treatment with negative control miRNA. This enrichment demonstrates that both the GEM-miR-15a and its targets, *CHK1* and *WEE1*, are loaded into Argonaute.

**Fig. 6.**
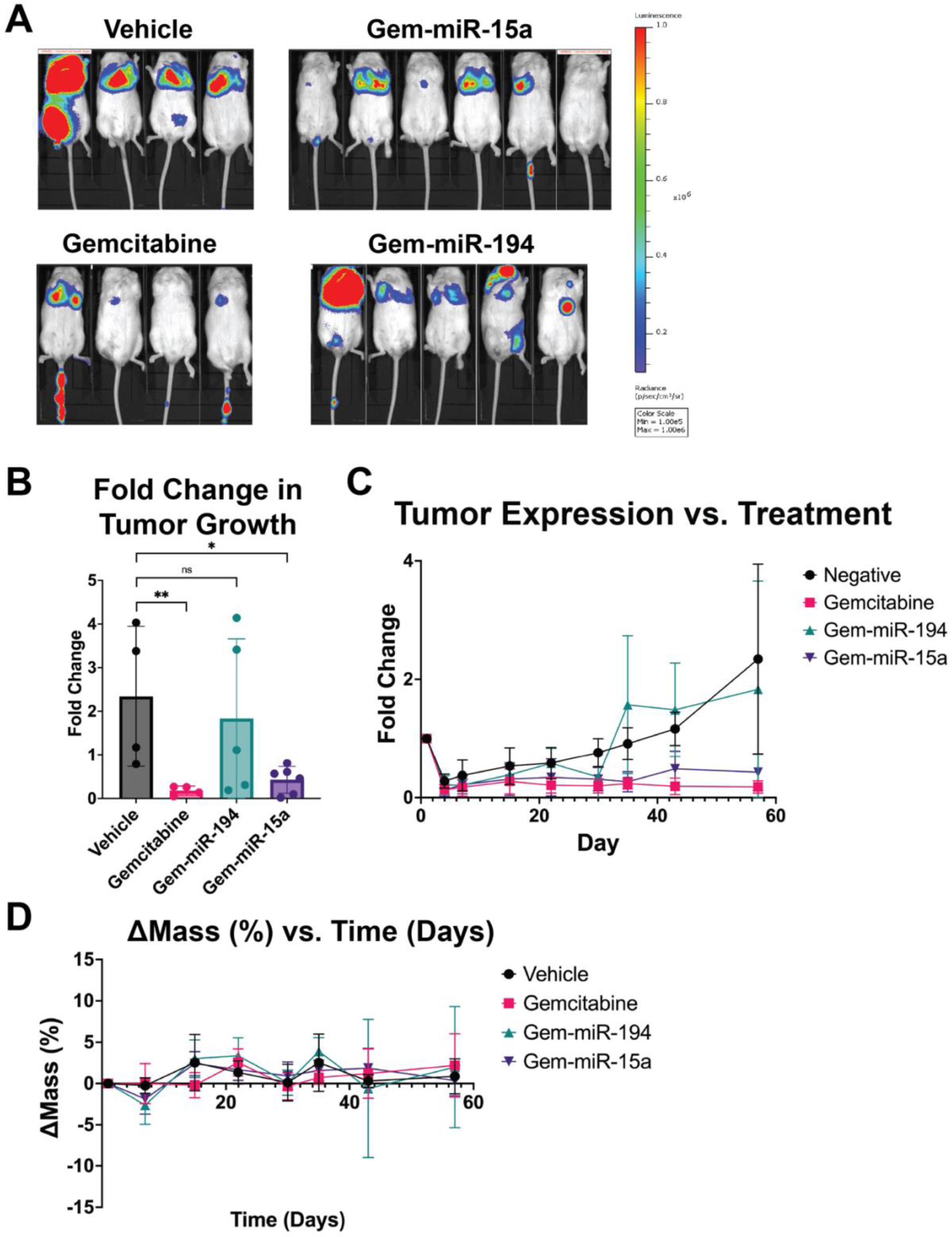
Gem-miR-15a, but not Gem-miR-194 is able to inhibit PDAC tumor growth *in vivo*. **(A)** Representative images of mice treated with vehicle control (Vehicle), Gem at 50 mg/kg (Gemcitabine), Gem-miR-194 at 4.0 mg/kg (Gem-miR-194), and Gem-miR-15a at 4.0 mg/kg (Gem-miR-15a) at day 57 (n = 4 - 6). Prior to treatment, xenografts of a luciferase-expressing PDAC cell line (Hs766T (+Luc)) were established in mice via intravenous (IV) tail vein injection. **(B)** By day 57, a significant reduction in PDAC tumor growth was only observed in mice treated with either Gem (*p* = 0.0083) or Gem-miR-15a (*p* = 0.0412), but not Gem-miR-194. **(C)** A timeline of PDAC tumor growth was made suggesting that Gem-miR-194 may initially reduce tumor growth before growing back to levels similar to the vehicle control group (Vehicle). **(D)** Body weight change was measured as an indicator of acute toxicity and no toxicity was observed (< 15% weight loss). Data are represented as mean ± SD. \**p* < 0.05 and \*\**p* < 0.01.

In summary, our results demonstrate the promising potential of modifying miRNAs with Gem as a strategy for developing miRNA-based therapeutics. Our results also demonstrate the potential for Gem-miR-15a as an anticancer treatment for patients with PDAC. Given the important regulatory function of miR-15a in immune response, other cancer types, and diseases, the therapeutic potential of Gem-miR-15a may extend beyond being an anticancer treatment for PDAC (*11, 37, 66–68*). As a general platform strategy, when utilizing this Gem modification on tumor suppressor miRNAs, the tumor suppressor miRNA candidate must be carefully selected based on their targets to reflect the complex nature of resistance in PDAC. However, modified miRNA-based therapies with purine and pyrimidine-based chemotherapeutic analogs can be conceptualized as a new class of multi-targeted biomimicry therapy with enhanced potency and minimal toxicity for future cancer care that can improve the quality of life for cancer patients.

## MATERIALS AND METHODS

### Study Design

Our study was designed with three main objectives. Our first objective was to examine the potency of Gem-miR-15a and Gem-miR-194 as a cancer therapeutic, with and without the use of a delivery vehicle, in PDAC 2D cell lines and patient-derived 3D organoids through dose-response assays. Gem-miR-15a and Gem-miR-194’s impact on cell cycle and apoptosis was also examined in PDAC 2D cell lines. Our second objective was to confirm whether our Gem-modified mimics retained miRNA function through western immunoblot analysis of reported targets of miR-15a and miR-194. We also performed AGO-CLIP with Gem-miR-15a to confirm our Gem-modified mimics can still be loaded into the RISC complex for RNAi. All *in vitro* experiments were performed with an n > 3. Finally, our third objective was to examine the potency of our Gem-modified mimics *in vivo* by comparing PDAC tumor growth via luciferase expression in mouse groups treated with either a vehicle control, Gem, Gem-miR-15a, or Gem-miR-194 with at least 4 mice per treatment group. Before treatment, mice were randomized into treatment groups based on average initial luciferase expression.

### Cell Lines and Tissue Culture

Human pancreatic cancer cell lines, AsPC-1, Hs766T, MIA PaCa-2, PANC-1, and Capan-1, were purchased from the American Type Culture Collection (ATCC, Manassas, VA, USA). The human immortalized pancreas epithelial cell line, hTERT-HPNE, was also purchased from the ATCC. Luciferase expressing Hs766T cells (Hs766T (+Luc)) were generated as previously described. AsPC-1 pancreatic cancer cells were cultured in RPMI-1640 Medium (Thermo Fisher Scientific, Waltham, MA, USA) supplemented with 10% fetal bovine serum (FBS) (MilliporeSigma, St. Louis, MO, USA). Hs766T, Hs766T (+Luc), MIA PaCA-2, and PANC-1 pancreatic cancer cells were cultured in Dulbecco’s Modified Eagle Medium (DMEM) (Thermo Fisher Scientific, Waltham, MA) supplemented with 10% FBS. Capan-1 cells were cultured in Iscove’s Modified Dulbecco’s medium (IMDM) (Thermo Fisher Scientific, Waltham, MA, USA) supplemented with 10% FBS. hTERT-HPNE cells were cultured in 75% DMEM without glucose (Thermo Fisher Scientific, Waltham, MA, USA) and 25% Medium M3 Base (Incell Corporation, San Antonio, TX, USA) supplemented with 5% FBS, human recombinant EGF (10 ng/ml) (Thermo Fisher Scientific, Waltham, MA, USA), D-glucose (5.5 mM) (MilliporeSigma, St. Louis, MO, USA), and puromycin (750 ng/ml) (MilliporeSigma, St. Louis, MO, USA).

### Design and Synthesis of Gemcitabine-Modified miRNA Mimics

Gem-modified miRNA mimics, Gem-miR-15a and Gem-miR-194, were designed and synthesized by substituting cytidine bases on the guide strands of hsa-miR-15a and hsa-miR-194-1 with Gem, respectively. The passenger strand was left unmodified to avoid any potential off-target effects and to preserve miRNA function. Oligonucleotides with these modifications as well as their corresponding passenger strand were purchased from Horizon Discovery (Horizon Discovery, Cambridge, UK). Both strands of oligonucleotides were HPLC purified. The guide strands and passenger strands were then annealed prior to use.

### miRNA Transfection

A non-specific scramble miRNA, Pre-miR^TM^ Negative Control #2 (negative control) (Thermo Fisher Scientific, Waltham, MA, USA) was used to represent the negative control. Pre-miR^TM^ hsa-miR-15a-5p and Pre-miR^TM^ hsa-miR-194-5p, miRNA mimics identical to the miR-15a-5p (miR-15a) and miR-194-5p (miR-194) strands, respectively, were purchased from Thermo Fisher Scientific (Thermo Fisher Scientific, Waltham, MA, USA). Cells were transfected with oligonucleotides at a concentration of 50 nM (negative control, miR-15a, Gem-miR-15a, and Gem-miR-194) or 200 nM (miR-194).

For vehicle-mediated transfections, cell lines were treated with Oligofectamine^TM^ (Thermo Fisher Scientific, Waltham, MA, USA) as previously described (*18, 39*). In brief, cell lines were seeded onto 6-well plates at a density of 100,00 cells/well. 24 hours later, cells were transfected with Oligofectamine^TM^ and their respective oligonucleotides according to the manufacturer’s protocol. Immediately after transfection, media was changed to fresh media supplemented with 10% dialyzed fetal bovine serum (DFBS) (Thermo Fisher Scientific, Waltham, MA, USA). For vehicle-free transfections, cell lines were seeded and treated as previously described. In brief, cell lines were seeded onto 96-well plates at a density of 1,000 cells/well. 24 hours after seeding, cells were transfected by changing media in the wells with fresh media mixed with their respective oligonucleotides. 24 hours after transfection, media was changed with fresh media supplemented with 10% DFBS.

### Cytotoxicity Analysis

For cytotoxicity analysis of oligonucleotides, cells were seeded and transfected as described above. For vehicle-mediated transfections, 24 hours post-transfection, cells were trypsinized and re-seeded onto a 96-well plate at a density of 1,000 cells/well. For vehicle-free transfections, cells were seeded onto 96-well plates at a density of 1,000 cells/well. Cells were transfected at varying concentrations of oligonucleotides for cytotoxicity. Cell viability was measured 6 days post-transfection using WST-1 Cell Proliferation Reagent (MilliporeSigma, St. Louis, MO, USA) according to the manufacturer’s protocol. In brief, cells were incubated with 10 μl of WST-1 per 100 μl of media for 1 hour at 37°C. After incubation, absorbance was measured at 450 and 630 nm using a SpectraMax i3 (Molecular Devices, San Jose, CA, USA) plate reader. The O.D. was calculated by subtracting the absorbance at 630 nm from that at 450 nm, and the relative proliferation was calculated by normalizing the O.D. to cells only transfected with vehicle. Absolute IC_50_ values were calculated using GraphPad Prism 9 (GraphPad Software, San Diego, CA, USA) using the following equation, where “Y” = 50.

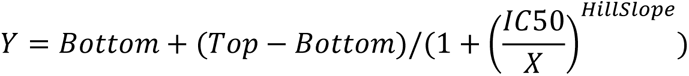

Cytoxicity analysis of Gem (MilliporeSigma, St. Louis, MO, USA) was measured by seeding cells onto a 96-well plate at 1,000 cells/well. 24 hours post-seeding, media was changed to fresh media supplemented with 10% DFBS and varying concentrations of Gem. 48 hours post-treatment, absorbance was measured at 450 and 630 nm using a SpectraMax i3 plate reader, and the absolute IC_50_ values were calculated as described above.

### Organoid Culture and Treatment

hF3, hF44, and hT89 organoid cells were provided by the Stony Brook Medicine Biobank (Stony Brook Medicine Biobank, Renaissance School of Medicine, Stony Brook, USA). Organoid cells were cultured and treated as previously described (*41*). In brief, organoid cells were cultured with Corning^TM^ Matrigel^TM^ GFR Membrane Matrix (Corning Incorporated, Corning, NY, USA) and incubated at 37°C in Human Complete Feeding Medium (hCPLT) consisting of the following: Advanced DMEM/F12 (Thermo Fisher Scientific, Waltham, MA, USA), HEPES (final concentration, 10 mmol/L) (Thermo Fisher Scientific, Waltham, MA, USA), GlutaMAX (final concentration, 1x) (Thermo Fisher Scientific, Waltham, MA, USA), A 83-01 (final concentration, 500 nmol/L) (Tocris Bioscience, Bristol, UK), hEGF (final concentration, 50 ng/mL) (Thermo Fisher Scientific, Waltham, MA, USA), mNoggin (final concentration, 100 ng/mL) (Thermo Fisher Scientific, Waltham, MA, USA), hFGF10 (final concentration, 100 ng/mL) (Thermo Fisher Scientific, Waltham, MA, USA), hGastrin I (final concentration, 0.01 μmol/L) (Tocris Bioscience, Bristol, UK), *N*-acetylcysteine (final concentration, 1.25 mmol/L) (MilliporeSigma, St. Louis, MO, USA), nicotinamide (final concentration, 10 mmol/L) (MilliporeSigma, St. Louis, MO, USA), prostaglandin E2 (final concentration, 1 μmol/L) (Tocris Bioscience, Bristol, UK) B27 supplement (final concentration, 1X) (Thermo Fisher Scientific, Waltham, MA, USA), R-spondin1-Conditioned Media (final concentration, 10%) (Thermo Fisher Scientific, Waltham, MA, USA), and afamin/Wnt3A conditioned media (Stony Brook Medicine Biobank, Renaissance School of Medicine, Stony Brook, USA). For treatment with oligonucleotides and Gem, the organoids were first dissociated into single cells. Viable cells were plated at 1,000 cells/well in 50 μL 10% Matrigel/hCPLT. 24 hours after plating and visual verification of organoid reformation, therapeutic compounds were added according to their respective conditions. Cell viability was assessed using CellTiter-Glo as per the manufacturer’s instruction (Promega, Madison, WI, USA) on a SpectraMax i3 plate reader either at 48 hours post treatment (Gem) or at 6 days post treatment (oligonucleotides). Cytotoxicities of different treatment conditions were then calculated as described above.

### Western Immunoblot Analysis

PDAC cells were seeded and transfected with or without a transfection vehicle as described above. Cells were transfected with their respective oligonucleotides and concentrations as described above. 72 hours post-transfection, cells were lysed with a mixture of RIPA buffer (MilliporeSigma, St. Louis, MO, USA) and protease inhibitor cocktail (MilliporeSigma, St. Louis, MO, USA), and the protein samples were the collected and used for western immunoblot analysis. Proteins were probed with rabbit anti-BMI1 antibody (Cell Signaling, 6964, 1:1000), mouse anti-CHK1 antibody (Cell Signaling, 2360, 1:1000), rabbit anti-FOXA1 antibody (Abcam, ab23738, 1:1000), rabbit anti-WEE1 antibody (Cell Signaling, 13084, 1:1000), rabbit anti-YAP1 antibody (Cell Signaling, 4912, 1:1000), or mouse anti-GAPDH antibody (Santa Cruz, sc47724, 1:100,000). Primary antibodies were diluted in 5% milk (Bio-Rad, Hercules, CA, USA) in TBST. After staining with primary antibodies, proteins were probed with either secondary antibodies goat anti-mouse-HRP (Bio-Rad, 1706516, 1:5000) or goat anti-rabbit-HRP (Bio-Rad, 1721019, 1:5000) depending on the used primary antibody. Protein bands were visualized using a LI-COR Biosciences Odyssey FC imaging system after the addition of SuperSignal™ West Pico PLUS Chemiluminescent Substrate (Thermo Fisher Scientific). Proteins were then quantified with Image Studio Version 5.2.4 (LI-COR Biosciences, Lincoln, NE, USA).

### AGO-CLIP

10 cm plates of PANC-1 cells were transfected with Oligofectamine^TM^ as described above. Cells were transfected with either negative control (50 nM) or Gem-miR-15a (50 nM). 24 hours later, the cells were irradiated with UV light and the immunoprecipitation of the crosslinked protein:RNA complexes were performed as previously described (*40*). Protein:RNA complexes were probed with mouse anti-pan Ago antibody (Millipore, MABE56, 1:1000) diluted in 5% milk in TBST. The protein:RNA complex was then stained with goat anti-mouse-HRP and visualized/quantified as described above.

### Cell Cycle Analysis

PDAC cells were seeded onto 6-well plates at a density of 100,000 cells/well. 24 hours after seeding plates, cells were washed with DPBS (Thermo Fisher Scientific, Waltham, MA, USA) and media was changed to fresh, serum-free media according to their respective lines to synchronize cells. PDAC cells were then transfected with their respective oligonucleotides and concentrations as described above or treated with 150 nM Gem (the concentration equivalent to Gem in Gem-miR-15a). 24 hours post-transfection, cells were resuspended in Krishan modified buffer supplemented with 0.02 mg/mL RNase H (Thermo Fisher Scientific, Waltham, MA, USA) and 0.05 mg/mL propidium iodide (MilliporeSigma, St. Louis, MO, USA). Cells were then analyzed by flow cytometry via CytoFLEX Flow Cytometer (Beckman Coulter, Brea, CA, USA) and results were analyzed by Modfit LT Software (BD Biosciences, Sparks, MD, USA).

### Apoptosis Assay

PDAC cells were plated and transfected with their respective treatment conditions via vehicle-mediated transfection as described above. 72 hours post-transfection, the cells were stained with Annexin V (Thermo Fisher Scientific, Waltham, MA, USA) and propidium iodide (MilliporeSigma, St. Louis, MO, USA). Apoptotic cells were quantified by flow cytometric analysis on a CytoFLEX Flow Cytometer (Beckman Coulter, Brea, CA, USA).

### Metastatic Pancreatic Cancer Mouse Model

All animal procedures were approved by the Stony Brook University Institutional Animal Care and Use Committee (IACUC) before the start of *in vivo* experiments. Non-obese diabetic (NOD)/severe combined immunodeficiency (SCID) mice (JAX: 001303) were purchased from The Jackson Laboratory (The Jackson Laboratory, Bar Harbor, ME, USA). 8-week-old NOD/SCID mice (2-3 males per group and 2-3 females per group) were inoculated with 2 x 10^6^ Hs766T (+Luc) cells suspended in 0.1 mL of PBS via intravenous (IV) tail vein injection. Mice were then divided into four groups (vehicle control, Gem-miR-15a, Gem-miR-194, and Gem). After confirming successful inoculation of Hs766T (+Luc) cells *in vivo*, two days post-inoculation, mice were treated with either 80 μg of PEI-MAX (Polysciences, Warrington, PA, USA) in 5% D-glucose (vehicle control group) (Thermo Fisher Scientific, Waltham, MA, USA), 80 µg (4.0 mg/kg) of Gem-miR-15a miRNA with 80 μg of PEI-MAX in 5% D-glucose (Gem-miR-15a group), 80 µg (4.0 mg/kg) of Gem-miR-194 miRNA with 80 μg of PEI-MAX in 5% D-glucose (Gem-miR-194 group), or 50 mg/kg of Gem in 1X DPBS (Gem group). Vehicle control, Gem-miR-15a, and Gem-miR-194 groups were treated on alternating days for 2 weeks for a total of 8 doses by IV tail vein injections. The Gem group was treated every 3 days for 2 weeks for a total of 4 doses by intraperitoneal (IP) injections. Using the same methods and conditions described above, *in vivo* experiments were repeated with the vehicle control group, Gem-miR-15a group, and the Gem group, with the concentration of Gem decreased to 12 mg/kg. Vehicle concentrations were selected in a non-toxic range as per manufacturer recommendations.

Luciferase expression was used to measure tumor growth via the IVIS Spectrum *In Vivo* Imaging System (IVIS) (PerkinElmer, Waltham, MA, USA) as previously described. In brief, mice were injected with IVISBrite D-Luciferin, RediJect (PerkinElmer, Waltham, MA, USA). 10 minutes post-injection, *in vivo* luciferase expression was measured via IVIS. Tumor growth was calculated as a function of (total flux at time of measurement [p/s])/(total flux at initial measurement [p/s]).

### Statistical Analysis

The quantitative data were presented as mean value ± standard deviation of at least 3 independent experiments in all studies unless otherwise specified. Statistical significance between two groups was determined using Student’s t-test. For comparisons with more than two groups, one-way ANOVA followed by a Tukey’s multiple comparisons test for pairwise comparisons were performed. For comparisons with non-parametric data with more than two groups, a Kruskal-Wallis followed by an uncorrected Dunn’s test was used. For comparisons with multiple variables, a two-way ANOVA followed by a Tukey’s multiple comparisons test were performed. A *p* value of less than 0.05 was considered statistically significant (\**p* < 0.05, \*\**p* < 0.01, \*\*\**p* < 0.001, \*\*\*\**p* < 0.0001).

## Acknowledgments

The authors wish to acknowledge the support of the Stony Brook Cancer Center Tissue Analytics Shared Resource for expert assistance with the development and propagation of human pancreatic cancer organoids.

## Funding

This research was supported by, in part, by the following:

USA National Institute of Health/National Cancer Institute (NIH/NCI), grant number R01CA1550197098 (JJ)

Curamir Therapeutics Inc. (JJ)

Veteran Affairs Merit Award, grant number BX005260-01 (JJ)

Stony Brook University INDUCER Award, grant number R25CA214272 (JJ)

## Author contributions

Conceptualization: JGY, GH, AF, JJ

Methodology: JGY, GH, AF, JJ

Investigation: JGY, GH, AF, EI, AP, AO

Visualization: JGY, GH

Funding acquisition: JJ

Project administration: JGY, GH, JJ

Supervision: JJ

Writing – original draft: JGY, GH

Writing – review & editing: JGY, GH, AF, JJ

## Data and materials availability

All data are available in the main text or the supplementary materials. Raw data supporting this study are available from the corresponding author (JJ) upon request.

## References and Notes

1. R. L. Siegel, K. D. Miller, N. S. Wagle, A. Jemal, Cancer statistics, 2023. CA Cancer J Clin 73, 17–48 (2023).

2. K. D. Miller, L. Nogueira, T. Devasia, A. B. Mariotto, K. R. Yabroff, A. Jemal, J. Kramer, R. L. Siegel, Cancer treatment and survivorship statistics, 2022. CA Cancer J Clin 72, 409–436 (2022).

3. Z. Wang, Y. Li, A. Ahmad, S. Banerjee, A. S. Azmi, D. Kong, F. H. Sarkar, Pancreatic cancer: understanding and overcoming chemoresistance. Nat Rev Gastroenterol Hepatol 8, 27–33 (2011).

4. P. E. Oberstein, K. P. Olive, Pancreatic cancer: why is it so hard to treat? Therap Adv Gastroenterol 6, 321–337 (2013).

5. P. Dauer, A. Nomura, A. Saluja, S. Banerjee, Microenvironment in determining chemo-resistance in pancreatic cancer: Neighborhood matters. Pancreatology 17, 7–12 (2017).

6. D. D. Von Hoff, T. Ervin, F. P. Arena, E. G. Chiorean, J. Infante, M. Moore, T. Seay, S. A. Tjulandin, W. W. Ma, M. N. Saleh, M. Harris, M. Reni, S. Dowden, D. Laheru, N. Bahary, R. K. Ramanathan, J. Tabernero, M. Hidalgo, D. Goldstein, E. Van Cutsem, X. Wei, J. Iglesias, M. F. Renschler, Increased survival in pancreatic cancer with nab-paclitaxel plus gemcitabine. N Engl J Med 369, 1691–1703 (2013).

7. T. Conroy, F. Desseigne, M. Ychou, O. Bouche, R. Guimbaud, Y. Becouarn, A. Adenis, J. L. Raoul, S. Gourgou-Bourgade, C. de la Fouchardiere, J. Bennouna, J. B. Bachet, F. Khemissa-Akouz, D. Pere-Verge, C. Delbaldo, E. Assenat, B. Chauffert, P. Michel, C. Montoto-Grillot, M. Ducreux, U. Groupe Tumeurs Digestives of, P. Intergroup, FOLFIRINOX versus gemcitabine for metastatic pancreatic cancer. N Engl J Med 364, 1817–1825 (2011).

8. T. Conroy, P. Hammel, M. Hebbar, M. Ben Abdelghani, A. C. Wei, J. L. Raoul, L. Chone, E. Francois, P. Artru, J. J. Biagi, T. Lecomte, E. Assenat, R. Faroux, M. Ychou, J. Volet, A. Sauvanet, G. Breysacher, F. Di Fiore, C. Cripps, P. Kavan, P. Texereau, K. Bouhier-Leporrier, F. Khemissa-Akouz, J. L. Legoux, B. Juzyna, S. Gourgou, C. J. O’Callaghan, C. Jouffroy-Zeller, P. Rat, D. Malka, F. Castan, J. B. Bachet, G. Canadian Cancer Trials, G. I. P. G. the Unicancer, FOLFIRINOX or Gemcitabine as Adjuvant Therapy for Pancreatic Cancer. N Engl J Med 379, 2395–2406 (2018).

9. H. Tong, Z. Fan, B. Liu, T. Lu, The benefits of modified FOLFIRINOX for advanced pancreatic cancer and its induced adverse events: a systematic review and meta-analysis. Sci Rep 8, 8666 (2018).

10. J. K. Lam, M. Y. Chow, Y. Zhang, S. W. Leung, siRNA Versus miRNA as Therapeutics for Gene Silencing. Mol Ther Nucleic Acids 4, e252 (2015).

11. G. A. Calin, C. D. Dumitru, M. Shimizu, R. Bichi, S. Zupo, E. Noch, H. Aldler, S. Rattan, M. Keating, K. Rai, L. Rassenti, T. Kipps, M. Negrini, F. Bullrich, C. M. Croce, Frequent deletions and down-regulation of micro-RNA genes miR15 and miR16 at 13q14 in chronic lymphocytic leukemia. Proc Natl Acad Sci U S A 99, 15524–15529 (2002).

12. R. Rupaimoole, F. J. Slack, MicroRNA therapeutics: towards a new era for the management of cancer and other diseases. Nat Rev Drug Discov 16, 203–222 (2017).

13. P. E. Blower, J. H. Chung, J. S. Verducci, S. Lin, J. K. Park, Z. Dai, C. G. Liu, T. D. Schmittgen, W. C. Reinhold, C. M. Croce, J. N. Weinstein, W. Sadee, MicroRNAs modulate the chemosensitivity of tumor cells. Mol Cancer Ther 7, 1–9 (2008).

14. A. Cimmino, G. A. Calin, M. Fabbri, M. V. Iorio, M. Ferracin, M. Shimizu, S. E. Wojcik, R. I. Aqeilan, S. Zupo, M. Dono, L. Rassenti, H. Alder, S. Volinia, C. G. Liu, T. J. Kipps, M. Negrini, C. M. Croce, miR-15 and miR-16 induce apoptosis by targeting BCL2. Proc Natl Acad Sci U S A 102, 13944–13949 (2005).

15. P. Trang, J. B. Weidhaas, F. J. Slack, MicroRNAs as potential cancer therapeutics. Oncogene 27 **Suppl 2**, S52–57 (2008).

16. J. Hanna, G. S. Hossain, J. Kocerha, The Potential for microRNA Therapeutics and Clinical Research. Front Genet 10, 478 (2019).

17. M. Ito, Y. Miyata, M. Okada, Current clinical trials with non-coding RNA-based therapeutics in malignant diseases: A systematic review. Transl Oncol 31, 101634 (2023).

18. S. Guo, A. Fesler, W. Huang, Y. Wang, J. Yang, X. Wang, Y. Zheng, G. R. Hwang, H. Wang, J. Ju, Functional Significance and Therapeutic Potential of miR-15a Mimic in Pancreatic Ductal Adenocarcinoma. Mol Ther Nucleic Acids 19, 228–239 (2020).

19. X. J. Zhang, H. Ye, C. W. Zeng, B. He, H. Zhang, Y. Q. Chen, Dysregulation of miR-15a and miR-214 in human pancreatic cancer. J Hematol Oncol 3, 46 (2010).

20. a. a. d. h. e. Cancer Genome Atlas Research Network. Electronic address, N. Cancer Genome Atlas Research, Integrated Genomic Characterization of Pancreatic Ductal Adenocarcinoma. Cancer Cell 32, 185–203 e113 (2017).

21. R. A. Moffitt, R. Marayati, E. L. Flate, K. E. Volmar, S. G. Loeza, K. A. Hoadley, N. U. Rashid, L. A. Williams, S. C. Eaton, A. H. Chung, J. K. Smyla, J. M. Anderson, H. J. Kim, D. J. Bentrem, M. S. Talamonti, C. A. Iacobuzio-Donahue, M. A. Hollingsworth, J. J. Yeh, Virtual microdissection identifies distinct tumor- and stroma-specific subtypes of pancreatic ductal adenocarcinoma. Nat Genet 47, 1168–1178 (2015).

22. M. Chan-Seng-Yue, J. C. Kim, G. W. Wilson, K. Ng, E. F. Figueroa, G. M. O’Kane, A. A. Connor, R. E. Denroche, R. C. Grant, J. McLeod, J. M. Wilson, G. H. Jang, A. Zhang, A. Dodd, S. B. Liang, A. Borgida, D. Chadwick, S. Kalimuthu, I. Lungu, J. M. S. Bartlett, P. M. Krzyzanowski, V. Sandhu, H. Tiriac, F. E. M. Froeling, J. M. Karasinska, J. T. Topham, D. J. Renouf, D. F. Schaeffer, S. J. M. Jones, M. A. Marra, J. Laskin, R. Chetty, L. D. Stein, G. Zogopoulos, B. Haibe-Kains, P. J. Campbell, D. A. Tuveson, J. J. Knox, S. E. Fischer, S. Gallinger, F. Notta, Transcription phenotypes of pancreatic cancer are driven by genomic events during tumor evolution. Nat Genet 52, 231–240 (2020).

23. C. H. Pan, Y. Otsuka, B. Sridharan, M. Woo, C. V. Leiton, S. Babu, M. Torrente Goncalves, R. R. Kawalerski, K. B. JD, D. K. Chang, A. V. Biankin, L. Scampavia, T. Spicer, L. F. Escobar-Hoyos, K. R. Shroyer, An unbiased high-throughput drug screen reveals a potential therapeutic vulnerability in the most lethal molecular subtype of pancreatic cancer. Mol Oncol 14, 1800–1816 (2020).

24. Z. Chang, Z. Li, X. Wang, Y. Kang, Y. Yuan, J. Niu, H. Wang, D. Chatterjee, J. B. Fleming, M. Li, J. L. Abbruzzese, P. J. Chiao, Deciphering the mechanisms of tumorigenesis in human pancreatic ductal epithelial cells. Clin Cancer Res 19, 549–559 (2013).

25. L. Zender, M. S. Spector, W. Xue, P. Flemming, C. Cordon-Cardo, J. Silke, S. T. Fan, J. M. Luk, M. Wigler, G. J. Hannon, D. Mu, R. Lucito, S. Powers, S. W. Lowe, Identification and validation of oncogenes in liver cancer using an integrative oncogenomic approach. Cell 125, 1253–1267 (2006).

26. A. Kapoor, W. Yao, H. Ying, S. Hua, A. Liewen, Q. Wang, Y. Zhong, C. J. Wu, A. Sadanandam, B. Hu, Q. Chang, G. C. Chu, R. Al-Khalil, S. Jiang, H. Xia, E. Fletcher-Sananikone, C. Lim, G. I. Horwitz, A. Viale, P. Pettazzoni, N. Sanchez, H. Wang, A. Protopopov, J. Zhang, T. Heffernan, R. L. Johnson, L. Chin, Y. A. Wang, G. Draetta, R. A. DePinho, Yap1 activation enables bypass of oncogenic Kras addiction in pancreatic cancer. Cell 158, 185–197 (2014).

27. B. Zhao, X. Wei, W. Li, R. S. Udan, Q. Yang, J. Kim, J. Xie, T. Ikenoue, J. Yu, L. Li, P. Zheng, K. Ye, A. Chinnaiyan, G. Halder, Z. C. Lai, K. L. Guan, Inactivation of YAP oncoprotein by the Hippo pathway is involved in cell contact inhibition and tissue growth control. Genes Dev 21, 2747–2761 (2007).

28. L. L. Parker, H. Piwnica-Worms, Inactivation of the p34cdc2-cyclin B complex by the human WEE1 tyrosine kinase. Science 257, 1955–1957 (1992).

29. B. Gabrielli, K. Brooks, S. Pavey, Defective cell cycle checkpoints as targets for anti-cancer therapies. Front Pharmacol 3, 9 (2012).

30. S. Leijen, R. M. van Geel, A. C. Pavlick, R. Tibes, L. Rosen, A. R. Razak, R. Lam, T. Demuth, S. Rose, M. A. Lee, T. Freshwater, S. Shumway, L. W. Liang, A. M. Oza, J. H. Schellens, G. I. Shapiro, Phase I Study Evaluating WEE1 Inhibitor AZD1775 As Monotherapy and in Combination With Gemcitabine, Cisplatin, or Carboplatin in Patients With Advanced Solid Tumors. J Clin Oncol 34, 4371–4380 (2016).

31. S. Leijen, R. M. van Geel, G. S. Sonke, D. de Jong, E. H. Rosenberg, S. Marchetti, D. Pluim, E. van Werkhoven, S. Rose, M. A. Lee, T. Freshwater, J. H. Beijnen, J. H. Schellens, Phase II Study of WEE1 Inhibitor AZD1775 Plus Carboplatin in Patients With TP53-Mutated Ovarian Cancer Refractory or Resistant to First-Line Therapy Within 3 Months. J Clin Oncol 34, 4354–4361 (2016).

32. P. Dong, M. Kaneuchi, H. Watari, J. Hamada, S. Sudo, J. Ju, N. Sakuragi, MicroRNA-194 inhibits epithelial to mesenchymal transition of endometrial cancer cells by targeting oncogene BMI-1. Mol Cancer 10, 99 (2011).

33. X. Zhu, D. Li, F. Yu, C. Jia, J. Xie, Y. Ma, S. Fan, H. Cai, Q. Luo, Z. Lv, L. Fan, miR-194 inhibits the proliferation, invasion, migration, and enhances the chemosensitivity of non-small cell lung cancer cells by targeting forkhead box A1 protein. Oncotarget 7, 13139–13152 (2016).

34. Z. Yuan, M. Ye, J. Qie, T. Ye, FOXA1 Promotes Cell Proliferation and Suppresses Apoptosis in HCC by Directly Regulating miR-212-3p/FOXA1/AGR2 Signaling Pathway. Onco Targets Ther 13, 5231–5240 (2020).

35. J. S. Roe, C. I. Hwang, T. D. D. Somerville, J. P. Milazzo, E. J. Lee, B. Da Silva, L. Maiorino, H. Tiriac, C. M. Young, K. Miyabayashi, D. Filippini, B. Creighton, R. A. Burkhart, J. M. Buscaglia, E. J. Kim, J. L. Grem, A. J. Lazenby, J. A. Grunkemeyer, M. A. Hollingsworth, P. M. Grandgenett, M. Egeblad, Y. Park, D. A. Tuveson, C. R. Vakoc, Enhancer Reprogramming Promotes Pancreatic Cancer Metastasis. Cell 170, 875–888 e820 (2017).

36. A. K. Beutel, C. J. Halbrook, Barriers and opportunities for gemcitabine in pancreatic cancer therapy. Am J Physiol Cell Physiol 324, C540–C552 (2023).

37. A. Fesler, H. Liu, J. Ju, Modified miR-15a has therapeutic potential for improving treatment of advanced stage colorectal cancer through inhibition of BCL2, BMI1, YAP1 and DCLK1. Oncotarget 9, 2367–2383 (2018).

38. N. Wu, A. Fesler, H. Liu, J. Ju, Development of novel miR-129 mimics with enhanced efficacy to eliminate chemoresistant colon cancer stem cells. Oncotarget 9, 8887–8897 (2018).

39. G. R. Hwang, J. G. Yuen, A. Fesler, H. Farley, J. D. Haley, J. Ju, Development of a 5-FU modified miR-129 mimic as a therapeutic for non-small cell lung cancer. Mol Ther Oncolytics 28, 277–292 (2023).

40. S. W. Chi, J. B. Zang, A. Mele, R. B. Darnell, Argonaute HITS-CLIP decodes microRNA-mRNA interaction maps. Nature 460, 479–486 (2009).

41. S. F. Boj, C. I. Hwang, L. A. Baker, Chio, II, D. D. Engle, V. Corbo, M. Jager, M. Ponz-Sarvise, H. Tiriac, M. S. Spector, A. Gracanin, T. Oni, K. H. Yu, R. van Boxtel, M. Huch, K. D. Rivera, J. P. Wilson, M. E. Feigin, D. Ohlund, A. Handly-Santana, C. M. Ardito-Abraham, M. Ludwig, E. Elyada, B. Alagesan, G. Biffi, G. N. Yordanov, B. Delcuze, B. Creighton, K. Wright, Y. Park, F. H. Morsink, I. Q. Molenaar, I. H. Borel Rinkes, E. Cuppen, Y. Hao, Y. Jin, I. J. Nijman, C. Iacobuzio-Donahue, S. D. Leach, D. J. Pappin, M. Hammell, D. S. Klimstra, O. Basturk, R. H. Hruban, G. J. Offerhaus, R. G. Vries, H. Clevers, D. A. Tuveson, Organoid models of human and mouse ductal pancreatic cancer. Cell 160, 324–338 (2015).

42. C. Wang, X. Li, L. Zhang, Y. Chen, R. Dong, J. Zhang, J. Zhao, X. Guo, G. Yang, Y. Li, C. Gu, Q. Xi, R. Zhang, miR-194-5p down-regulates tumor cell PD-L1 expression and promotes anti-tumor immunity in pancreatic cancer. Int Immunopharmacol 97, 107822 (2021).

43. J. G. Yuen, A. Fesler, G. R. Hwang, L. B. Chen, J. Ju, Development of 5-FU-modified tumor suppressor microRNAs as a platform for novel microRNA-based cancer therapeutics. Mol Ther 30, 3450–3461 (2022).

44. Y. H. Soung, H. Chung, C. Yan, A. Fesler, H. Kim, E. S. Oh, J. Ju, J. Chung, Therapeutic Potential of Chemically Modified miR-489 in Triple-Negative Breast Cancers. Cancers (Basel) 12, (2020).

45. E. Driehuis, K. Kretzschmar, H. Clevers, Establishment of patient-derived cancer organoids for drug-screening applications. Nat Protoc 15, 3380–3409 (2020).

46. M. Huch, J. A. Knoblich, M. P. Lutolf, A. Martinez-Arias, The hope and the hype of organoid research. Development 144, 938–941 (2017).

47. U. Wellner, J. Schubert, U. C. Burk, O. Schmalhofer, F. Zhu, A. Sonntag, B. Waldvogel, C. Vannier, D. Darling, A. zur Hausen, V. G. Brunton, J. Morton, O. Sansom, J. Schuler, M. P. Stemmler, C. Herzberger, U. Hopt, T. Keck, S. Brabletz, T. Brabletz, The EMT-activator ZEB1 promotes tumorigenicity by repressing stemness-inhibiting microRNAs. Nat Cell Biol 11, 1487–1495 (2009).

48. K. Schlegelmilch, M. Mohseni, O. Kirak, J. Pruszak, J. R. Rodriguez, D. Zhou, B. T. Kreger, V. Vasioukhin, J. Avruch, T. R. Brummelkamp, F. D. Camargo, Yap1 acts downstream of alpha-catenin to control epidermal proliferation. Cell 144, 782–795 (2011).

49. M. Overholtzer, J. Zhang, G. A. Smolen, B. Muir, W. Li, D. C. Sgroi, C. X. Deng, J. S. Brugge, D. A. Haber, Transforming properties of YAP, a candidate oncogene on the chromosome 11q22 amplicon. Proc Natl Acad Sci U S A 103, 12405–12410 (2006).

50. L. A. Fernandez, M. Squatrito, P. Northcott, A. Awan, E. C. Holland, M. D. Taylor, Z. Nahle, A. M. Kenney, Oncogenic YAP promotes radioresistance and genomic instability in medulloblastoma through IGF2-mediated Akt activation. Oncogene 31, 1923–1937 (2012).

51. L. A. Parsels, J. D. Parsels, D. M. Tanska, J. Maybaum, T. S. Lawrence, M. A. Morgan, The contribution of DNA replication stress marked by high-intensity, pan-nuclear gammaH2AX staining to chemosensitization by CHK1 and WEE1 inhibitors. Cell Cycle 17, 1076–1086 (2018).

52. S. Chung, P. Vail, A. K. Witkiewicz, E. S. Knudsen, Coordinately Targeting Cell-Cycle Checkpoint Functions in Integrated Models of Pancreatic Cancer. Clin Cancer Res 25, 2290–2304 (2019).

53. H. Zhai, M. Karaayvaz, P. Dong, N. Sakuragi, J. Ju, Prognostic significance of miR-194 in endometrial cancer. Biomark Res 1, 12 (2013).

54. P. Huang, L. Xia, Q. Guo, C. Huang, Z. Wang, Y. Huang, S. Qin, W. Leng, D. Li, Genome-wide association studies identify miRNA-194 as a prognostic biomarker for gastrointestinal cancer by targeting ATP6V1F, PPP1R14B, BTF3L4 and SLC7A5. Front Oncol 12, 1025594 (2022).

55. M. Castaneda, P. D. Hollander, S. A. Mani, Forkhead Box Transcription Factors: Double-Edged Swords in Cancer. Cancer Res 82, 2057–2065 (2022).

56. B. Sahu, M. Laakso, K. Ovaska, T. Mirtti, J. Lundin, A. Rannikko, A. Sankila, J. P. Turunen, M. Lundin, J. Konsti, T. Vesterinen, S. Nordling, O. Kallioniemi, S. Hautaniemi, O. A. Janne, Dual role of FoxA1 in androgen receptor binding to chromatin, androgen signalling and prostate cancer. EMBO J 30, 3962–3976 (2011).

57. V. Kulkarni, A. R. Naqvi, J. R. Uttamani, S. Nares, MiRNA-Target Interaction Reveals Cell-Specific Post-Transcriptional Regulation in Mammalian Cell Lines. Int J Mol Sci 17, (2016).

58. S. Purser, P. R. Moore, S. Swallow, V. Gouverneur, Fluorine in medicinal chemistry. Chem Soc Rev 37, 320–330 (2008).

59. Y. Song, M. K. Washington, H. C. Crawford, Loss of FOXA1/2 is essential for the epithelial-to-mesenchymal transition in pancreatic cancer. Cancer Res 70, 2115–2125 (2010).

60. H. J. Jin, J. C. Zhao, I. Ogden, R. C. Bergan, J. Yu, Androgen receptor-independent function of FoxA1 in prostate cancer metastasis. Cancer Res 73, 3725–3736 (2013).

61. M. Teng, S. Zhou, C. Cai, M. Lupien, H. H. He, Pioneer of prostate cancer: past, present and the future of FOXA1. Protein Cell 12, 29–38 (2021).

62. N. Yamaguchi, Y. Nakayama, N. Yamaguchi, Down-regulation of Forkhead box protein A1 (FOXA1) leads to cancer stem cell-like properties in tamoxifen-resistant breast cancer cells through induction of interleukin-6. J Biol Chem 292, 8136–8148 (2017).

63. A. Parolia, M. Cieslik, S. C. Chu, L. Xiao, T. Ouchi, Y. Zhang, X. Wang, P. Vats, X. Cao, S. Pitchiaya, F. Su, R. Wang, F. Y. Feng, Y. M. Wu, R. J. Lonigro, D. R. Robinson, A. M. Chinnaiyan, Distinct structural classes of activating FOXA1 alterations in advanced prostate cancer. Nature 571, 413–418 (2019).

64. B. Song, S. H. Park, J. C. Zhao, K. W. Fong, S. Li, Y. Lee, Y. A. Yang, S. Sridhar, X. Lu, S. A. Abdulkadir, R. L. Vessella, C. Morrissey, T. M. Kuzel, W. Catalona, X. Yang, J. Yu, Targeting FOXA1-mediated repression of TGF-beta signaling suppresses castration-resistant prostate cancer progression. J Clin Invest 129, 569–582 (2019).

65. M. J. Jiang, Y. Y. Chen, J. J. Dai, D. N. Gu, Z. Mei, F. R. Liu, Q. Huang, L. Tian, Dying tumor cell-derived exosomal miR-194-5p potentiates survival and repopulation of tumor repopulating cells upon radiotherapy in pancreatic cancer. Mol Cancer 19, 68 (2020).

66. N. Liu, C. W. Chang, C. J. Steer, X. W. Wang, G. Song, MicroRNA-15a/16-1 Prevents Hepatocellular Carcinoma by Disrupting the Communication Between Kupffer Cells and Regulatory T Cells. Gastroenterology 162, 575–589 (2022).

67. V. M. D. Almanzar, K. Shah, J. F. LaComb, A. Mojumdar, H. R. Patel, J. Cheung, M. Tang, J. Ju, A. B. Bialkowska, 5-FU-miR-15a Inhibits Activation of Pancreatic Stellate Cells by Reducing YAP1 and BCL-2 Levels In Vitro. Int J Mol Sci 24, (2023).

68. J. D. Gagnon, R. Kageyama, H. M. Shehata, M. S. Fassett, D. J. Mar, E. J. Wigton, K. Johansson, A. J. Litterman, P. Odorizzi, D. Simeonov, B. J. Laidlaw, M. Panduro, S. Patel, L. T. Jeker, M. E. Feeney, M. T. McManus, A. Marson, M. Matloubian, S. Sanjabi, K. M. Ansel, miR-15/16 Restrain Memory T Cell Differentiation, Cell Cycle, and Survival. Cell Rep 28, 2169–2181 e2164 (2019).

