## Supplementary Figures for "Development of Gemcitabine-Modified miRNA Mimics as Cancer Therapeutics for Pancreatic Ductal Adenocarcinoma"

### List of Supplementary Figures

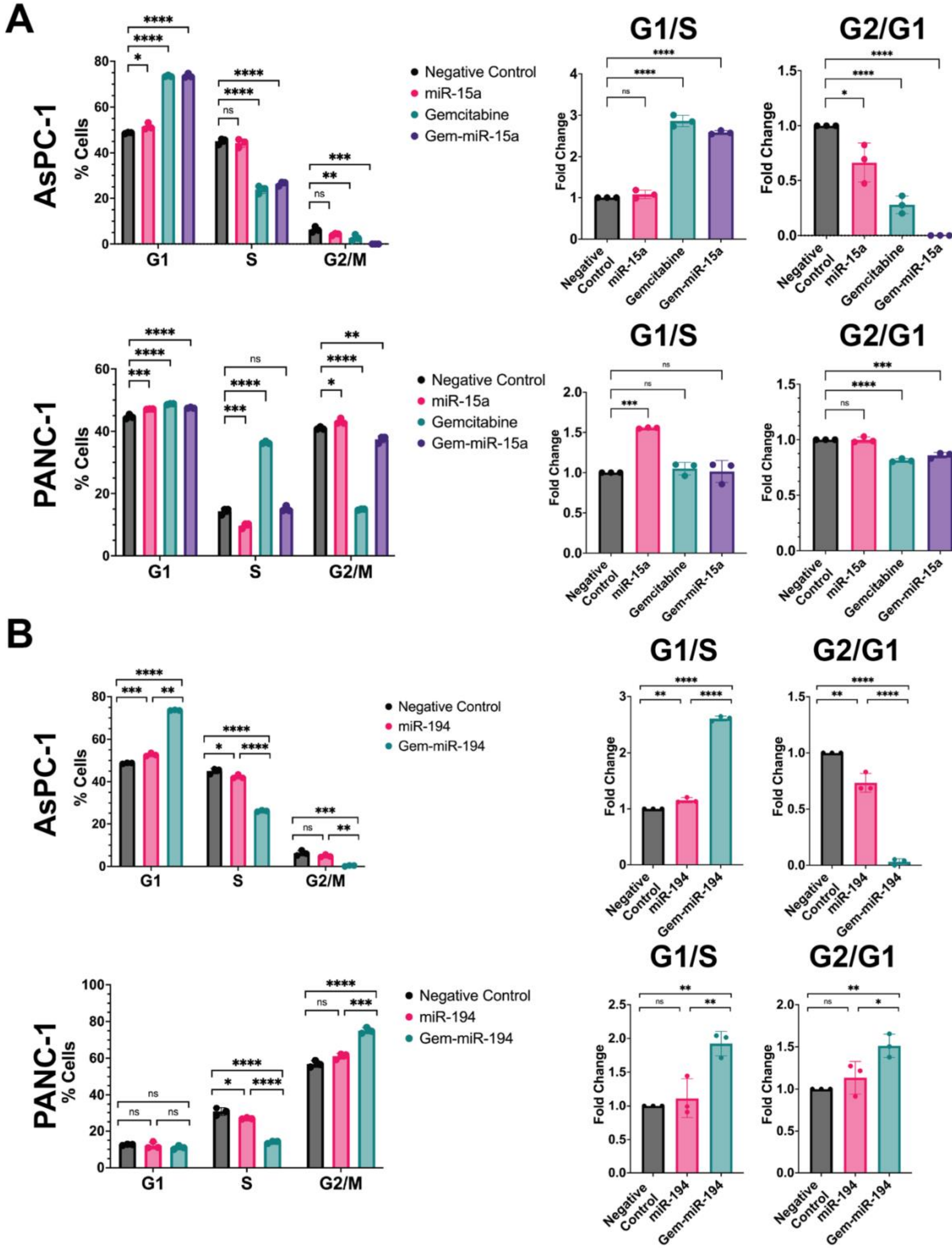

**Fig. S1: Gem-miR-15a and Gem-miR-194 induce cell cycle arrest in PDAC cell lines AsPC-1 and PANC-1. (A)** In AsPC-1 cells, Gem-miR-15a was found to induce cell cycle arrest by inducing an increase of cells in G1 ($p < 0.0001$) and a decrease of cells in S ($p < 0.0001$) and G2/M ($p = 0.0002$) (n = 3). An increase in G1/S ratio ($p < 0.0001$) and a decrease in G2/G1 ratio ($p < 0.0001$) was also observed, suggesting G1 phase arrest. In PANC-1 cells, Gem-miR-15a was found to induce an increase of cells in G1 ($p < 0.0001$) and a decrease of cells in G2 ($p = 0.0002$) (n = 3). A significant change in G1/S ratio was not observed, but a decrease in G2/G1 ratio ($p = 0.0001$) was observed, suggesting G1 phase arrest. **(B)** In AsPC-1 cells, Gem-miR-194 was also found to induce cell cycle arrest by inducing an increase of cells in G1 ($p < 0.0001$) and a decrease of cells in S ($p < 0.0001$) and G2/M ($p = 0.0003$) (n =3). An increase in G1/S ratio ($p < 0.0001$) and a decrease in G2/G1 ratio ($p < 0.0001$) was also observed, suggesting G1 phase arrest. In PANC-1 cells, Gem-miR-194 was found to induce a decrease of cells in S ($p < 0.0001$) and an increase of cells in G2/M ($p < 0.0001$) (n = 3). An increase in G1/S ratio ($p < 0.0030$) and G2/G1 ratio ($p < 0.0090$) was observed, suggesting that Gem-miR-194 induces either G1 or G2/M phase arrest. Data are represented as mean ± SD. *$p < 0.05$, **$p < 0.01$, ***$p < 0.001$, ****$p < 0.0001$.

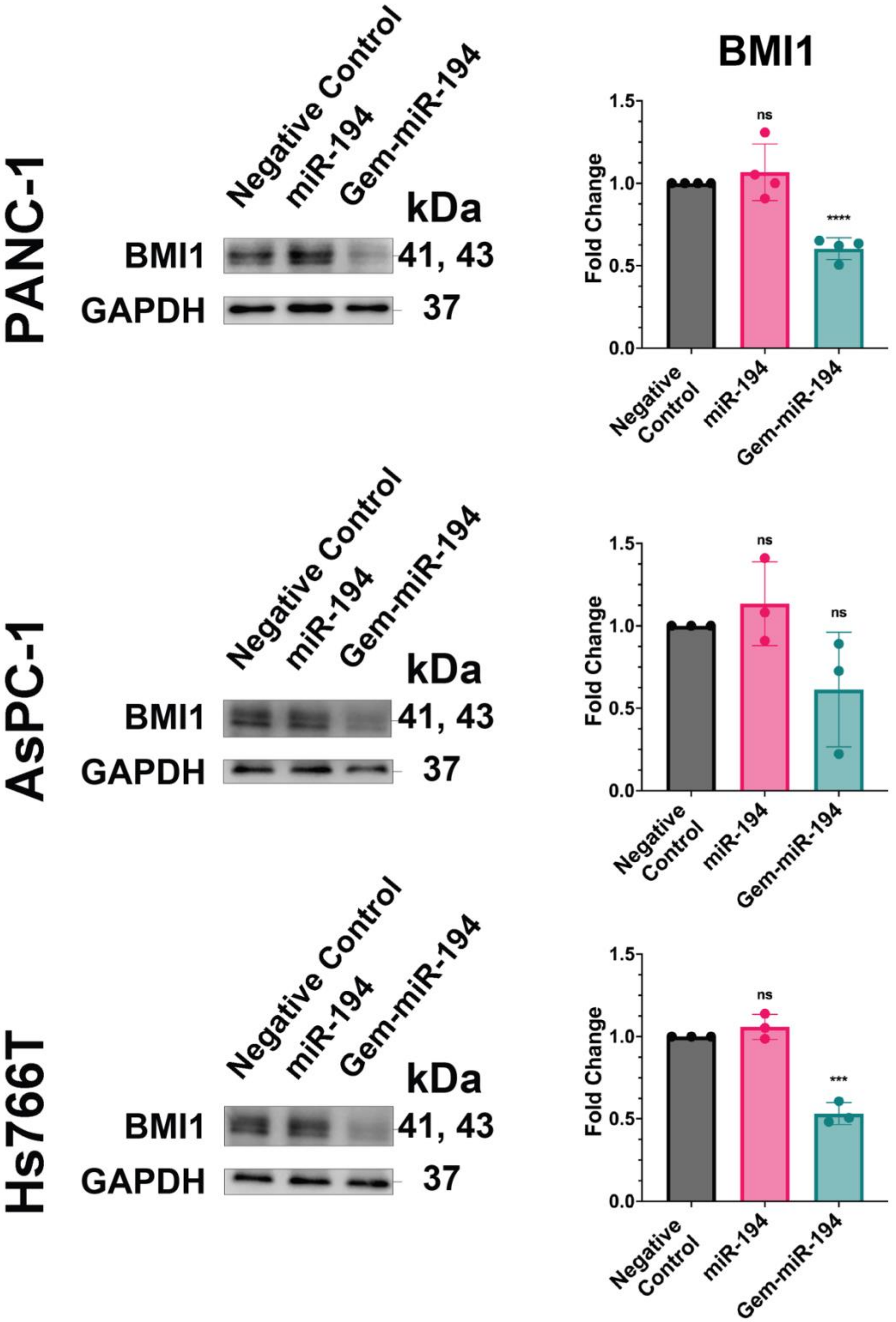
BMI1
PANC-1
AsPC-1
Hs766T
Negative Control
miR-194
Gem-miR-194
kDa
BMI1
GAPDH
41, 43
37
Fold Change
1.5
1.0
0.5
0.0
ns
****
***
Negative Control

**Fig. S2: Gem-miR-194 downregulates expression of BMI1 in PDAC in a cell-line dependent manner.** Gem-miR-194 was found to downregulate expression of miR-194 target, BMI1, only in PDAC cells PANC-1 ($p < 0.0001$, n = 4) and Hs766T ($p = 0.0003$, n = 3). In addition, despite being a known target of miR-194 in other cancer types, miR-194 was not found to downregulate expression of BMI1 in these PDAC cell lines. Data are represented as mean ± SD. ***$p < 0.001$, ****$p < 0.0001$.

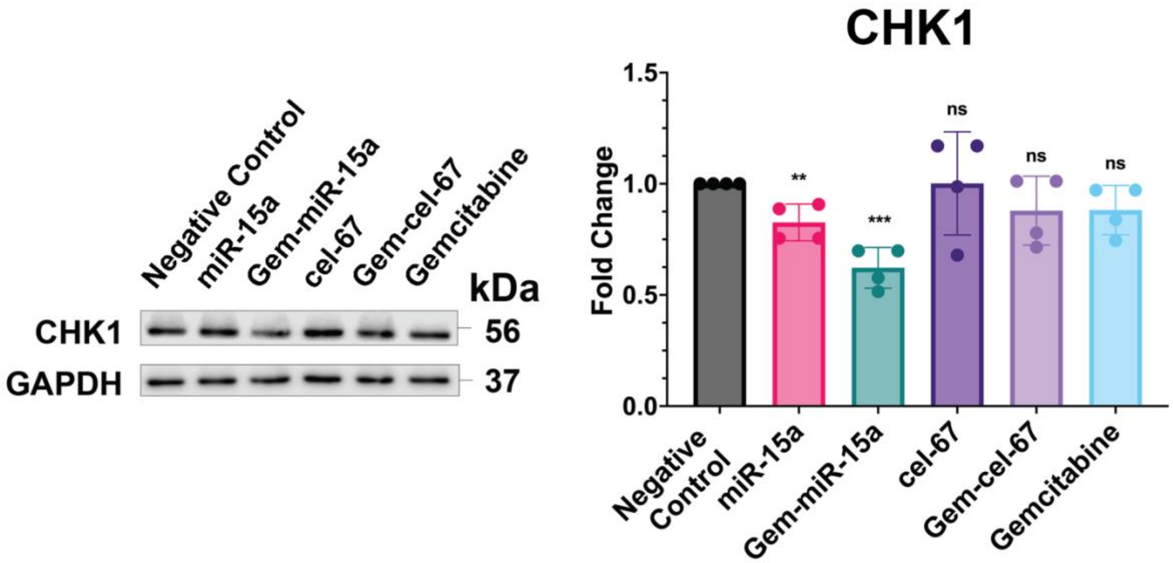


**Fig. S3: Gem-modified nonspecific miRNA mimic does not downregulate expression of miRNA target.** Using a nonspecific miRNA (cel-miR-67) sequence, a Gem-modified mimic (Gem-cel-67) was developed. Gem-cel-67 was found to not have a significant impact on miR-15a target, CHK1, compared to miR-15a and Gem-miR-15a, suggesting that modifying a miRNA sequence with Gem does not impart non-specific miRNA target specificity (n = 4). Data are represented as mean ± SD. **$p < 0.01$, ***$p < 0.001$.

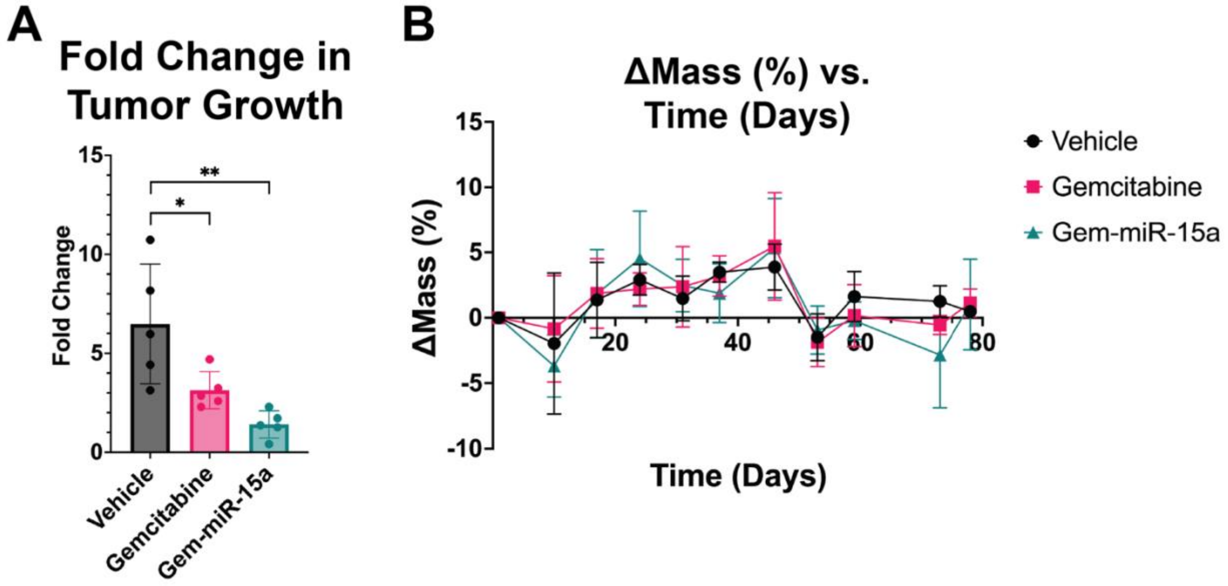


**Fig. S4: Gem-miR-15a inhibits PDAC tumor growth *in vivo*. (A)** The *in vivo* experiment shown in Fig. 6 was repeated with the following treatment groups: vehicle control (Vehicle), Gem (Gemcitabine, 12 mg/kg), and Gem-miR-15a (Gem-miR-15a, 4.0 mg/kg) (n = 5). By day 73, a significant reduction in PDAC tumor growth was observed in mice treated with Gem-miR-15a ($p = 0.0028$). **(B** Body weight change was measured as an indicator of acute toxicity and no toxicity was observed (< 15% weight loss). Data are represented as mean ± SD. *$p < 0.05$ and **$p < 0.01$.**)**
